# The RNA-binding ATPase, Armitage, Couples piRNA Amplification in Nuage to Phased piRNA Production on Mitochondria

**DOI:** 10.1101/445825

**Authors:** Daniel Tianfang Ge, Wei Wang, Cindy Tipping, Ildar Gainetdinov, Zhiping Weng, Phillip D. Zamore

## Abstract

PlWI-interacting RNAs (piRNAs) silence transposons in *Drosophila* ovaries, ensuring female fertility. Two coupled pathways generate germline piRNAs: the ping-pong cycle, in which the PIWI proteins Aubergine and Ago3 increase the abundance of pre-existing piRNAs, and the phased piRNA pathway, which generates strings of tail-to-head piRNAs, one after another. Proteins acting in the ping-pong cycle localize to nuage, whereas phased piRNA production requires Zucchini, an endonuclease on the mitochondrial surface. Here, we report that Armitage (Armi), an RNA-binding ATPase localized to both nuage and mitochondria, links the ping-pong cycle to the phased piRNA pathway. Mutations that block phased piRNA production deplete Armi from nuage. Armi ATPase mutants cannot support phased piRNA production and inappropriately bind mRNA instead of piRNA precursors. We propose that Armi shuttles between nuage and mitochondria, feeding precursor piRNAs generated by Ago3 cleavage into the Zucchini-dependent production of Aubergine- and Piwi-bound piRNAs on the mitochondrial surface.

## INTRODUCTION

In animals, PIWI-interacting RNAs (piRNAs) direct PIWI-clade Argonaute proteins to silence germline transposons, ensuring fertility (Czech and Hannon, 2016; Huang et al., 2017). In *Drosophila*, the cytoplasmic PIWI proteins Aubergine (Aub) and Ago3 increase piRNA abundance via reciprocal cleavage of sense transposon mRNAs and antisense piRNA precursor transcripts, a process termed the ping-pong amplification cycle (Brennecke et al., 2007; Gunawardane et al., 2007). The “initiator” and “responder” piRNAs amplified by the ping-pong cycle go on to direct production of tail-to-head strings of “trailing” phased piRNAs (Han et al., 2015a; Mohn et al., 2015). Typically, Ago3-catalyzed, piRNA-directed cleavage of piRNA precursor transcripts initiates the production of these phased piRNAs, which are then loaded into Piwi, and, to a lesser extent, Aub (Han et al., 2015a; Mohn et al., 2015; Wang et al., 2015). piRNA-bound Piwi, but not unloaded Piwi, can then transit to the nucleus, where it directs Histone 3 lysine 9 trimethylation (H3K9me3) to transposons, silencing transcription by generating heterochromatin (McCue and Slotkin, 2012; Le Thomas et al., 2013; Rozhkov et al., 2013; Yashiro et al., 2018). (Historically, the piRNAs generated by PIWI-catalyzed precursor cleavage were called “secondary” piRNAs, whereas the piRNAs now known to be phased have been termed “primary” piRNAs. Here, we use the terms *initiator* and *responder* for ping-pong piRNAs and *trailing* for phased piRNAs.)

Trailing piRNA production requires Zucchini (Zuc), an endonuclease proposed to simultaneously generate the 3’ end of the preceding immature piRNA (pre-piRNA) and the 5’ end of the precursor (pre-pre-piRNA) that will produce the next pre-piRNA (Han et al., 2015a; Mohn et al., 2015). Zuc belongs to the phospholipase D superfamily, whose HKD catalytic domain hydrolyzes phosphodiester bonds in phospholipids or nucleic acids (Pane et al., 2007; Selvy et al., 2011). Zuc cleaves single-stranded RNA in vitro, but without the NpU preference expected for an endonuclease generating phased piRNAs (Ipsaro et al., 2012; Nishimasu et al., 2012; Han et al., 2015a; Mohn et al., 2015; Nishida et al., 2018).

Phased piRNA production also requires Armitage (Armi), a member of the Upf1 family of ATP-dependent 5’-to-3’ helicase proteins (Cook et al., 2004; Haase et al., 2010; Olivieri et al., 2010; Saito et al., 2010). Strikingly, artificially tethering Armi to a transcript triggers its conversion into piRNAs (Pandey et al., 2017; Rogers et al., 2017). Armi is dispensable for the ping-pong cycle (Malone et al., 2009; Handler et al., 2011). The mammalian Armi homolog Mov10l1 binds RNA (Vourekas et al., 2015), and recombinant fly Armi unwinds RNA duplexes 5’-to-3’ (Pandey et al., 2017), suggesting that, like Upf1, Armi uses ATP to gate its RNA binding.

Loss of Zuc, Armi, or other phased-piRNA biogenesis proteins such as Piwi, Gasz (Germ cell protein with Ankyrin repeats, Sterile alpha motif, and leucine Zipper), Minotaur, and Papi, has little effect on ping-pong amplification (Handler et al., 2011; Baena-Lopez et al., 2013; Handler et al., 2013; Vagin et al., 2013; Han et al., 2015a; Hayashi et al., 2016). In contrast, germline phased-piRNA production collapses without the ping-pong machinery, because ping-pong amplification generates the initiator piRNAs that generate the pre-pre-RNAs that feed phased piRNA production (Han et al., 2015a; Mohn et al., 2015). The current model for piRNA biogenesis posits that phased-piRNA production begins when Ago3 cuts a complementary RNA, generating a 5’ monophosphorylated pre-pre-piRNA that can bind Aub. Next, Aub directs an endonuclease-likely Zuc-to cut the pre-pre-piRNA 5’ to the first downstream accessible uridine residue, simultaneously releasing Aub bound to a pre-piRNA and creating a new 5’ monophosphorylated pre-pre-piRNA from which trailing piRNAs can be generated (Mohn et al., 2015; Gainetdinov et al., 2018). Typically, Piwi binds the 5’ end of the new pre-pre-piRNA. Piwi directs Zuc to cut the pre-pre-piRNA 5’ to the next accessible uridine, generating a Piwi-bound, trailing pre-piRNA. The 3’ endonuclease cleavage product can then bind a new Piwi protein, repeating the process to generate another trailing piRNA and pre-pre-piRNA. The production of phased piRNAs is believed to be limited by the frequency of ping-pong piRNA-directed cleavages: more frequent cleavage yields shorter substrates for phased piRNA production (Mohn et al., 2015).

The initial products of both the ping-pong and phased piRNA pathways are pre-piRNAs bound to PIWI proteins. In flies, the 3’ ends of pre-piRNAs are subsequently trimmed by Nibbler and 2’-*O*-methylated by the S-adenosyl methionine-dependent methyl transferase, Hen1, completing piRNA biogenesis (Horwich et al., 2007; Saito et al., 2007; Feltzin et al., 2015; Hayashi et al., 2016; Wang et al., 2016). Unlike pre-piRNAs in many animals, fly pre-piRNAs are nearly the same length as mature piRNAs, so that many piRNAs, particularly Piwi-bound piRNAs, are often 2’-*O*-methylated without 3’ trimming (Hayashi et al., 2016; Gainetdinov et al., 2018).

What couples the ping-pong cycle to the phased piRNAs pathway? In *Drosophila* germline nurse cells, ping-pong piRNAs are thought to be made in perinuclear nuage—electron-dense, non-membrane-bound bodies (Mahowald, 1971; Eddy, 1975)-which contains the piRNA precursor transcripts and proteins required for ping-pong amplification (Liang et al., 1994; Brennecke et al., 2007; Lim and Kai, 2007; Nishida et al., 2009; Patil and Kai, 2010; Handler et al., 2011; Zhang et al., 2011; Olivieri et al., 2012; White-Cooper, 2012; Zhang et al., 2012a; Mohn et al., 2014; Patil et al., 2014; Andress et al., 2016). In contrast, trailing phased piRNAs are likely made on the mitochondrial outer membrane, where phasing factors are found (Choi et al., 2006; Olivieri et al., 2010; Saito et al., 2010; Schirle and MacRae, 2012; Baena-Lopez et al., 2013; Handler et al., 2013; Vagin et al., 2013; Huang et al., 2014; Izumi et al., 2016; Nishida et al., 2018). Here, we report that Armitage (Armi) localizes to both nuage and mitochondria and couples the ping-pong cycle to the phased piRNA pathway. Mutations that block phased piRNA production trap Armi on mitochondria. Armi ATPase mutants do not support phased piRNA production and cause Armi to inappropriately bind mRNA instead of piRNA precursors. We propose that Armi shuttles between nuage and mitochondria, feeding precursor piRNAs generated by Ago3 and Aub cleavage into the Zucchini-dependent production of Aubergine- and Piwi-bound piRNAs on the mitochondrial surface.

## RESULTS

### Nuage and Mitochondria are Physically Separate in Nurse Cells

In principle, physical association of nuage and mitochondria could explain the coupling of ping-pong amplification to phased piRNA production. Although nuage often associates with mitochondria (Eddy, 1975; Kloc et al., 2014), particularly in mammals where a nuage-like “cement” resides in the interstices of clustered mitochondria (Eddy, 1974), electron microscopy of *Drosophila* nurse cells rarely finds nuage apposed to mitochondria (Dapples and King, 1970; Mahowald, 1970; Mahowald, 1971; Liang et al., 1994; Wilsch-Bräuninger et al., 1997; Jaglarz et al., 2011). Quantitative immunofluorescence microscopy further confirms that most mitochondria do not contact nuage (Figure 1A, top row). In stage 3 egg chambers, the 15 germline nurse cell nuclei are well separated, allowing unambiguous detection of cytoplasmic proteins (Figure S1A); immunostaining of these cells readily detected the piRNA biogenesis proteins Aub, a representative ping-pong factor, and Zuc, a representative phased piRNA pathway component. The germline-specific DEAD box protein Vasa served as a marker for nuage, while ATP synthase complex V alpha subunit (ATP5A) identified the inner mitochondrial membrane. In wild-type ovaries, Vasa-containing nuage puncta surround the nurse cell nuclei. Mitochondria are evenly distributed in the cytoplasm, with a tendency to clump near the nucleus (Figure 1A, top row). Quantification of the fluorescent signal detected little Vasa coincident with mitochondria: 18% ± 9% of Vasa signal overlapped with ATP5A, while 88% ± 7% overlapped with Aub (*n* = 54 images from three egg chambers). Because the wavelength of the emitted light limits the resolution of fluorescence microscopy (λ/[2 × numerical aperture] where λ_emitted_ = 525 nm for Alexa Fluor 488), the limit of resolution for our experiments, ~188 nm, is not significantly smaller than the typical nuage puncta (Jaglarz et al., 2011). Consequently, fluorescence microscopy cannot distinguish between the biological finding of few mitochondria associating with nuage and artifacts caused by diffracted light.

To further test our observation that few mitochondria abut nuage, we used transmission electron microscopy (TEM), a technique that readily distinguishes between nuage, electron-dense, fibrous granules lacking a membrane, and mitochondria, double-membrane-bound organelles with characteristic internal cristae. In TEM images of stage 1 to stage 8 egg chambers, nuage and mitochondria rarely touched: in two independent experiments, none of 31 and two of 191 nuage puncta contacted mitochondria (Figure 1B and S1B). We conclude that *Drosophila* nurse cell nuage and mitochondria are physically separate, suggesting that one or more piRNA pathway proteins help escort the precursor RNAs generated by the ping-pong cycle from nuage to the mitochondrial surface for processing into phased piRNAs.

**Figure 1.**
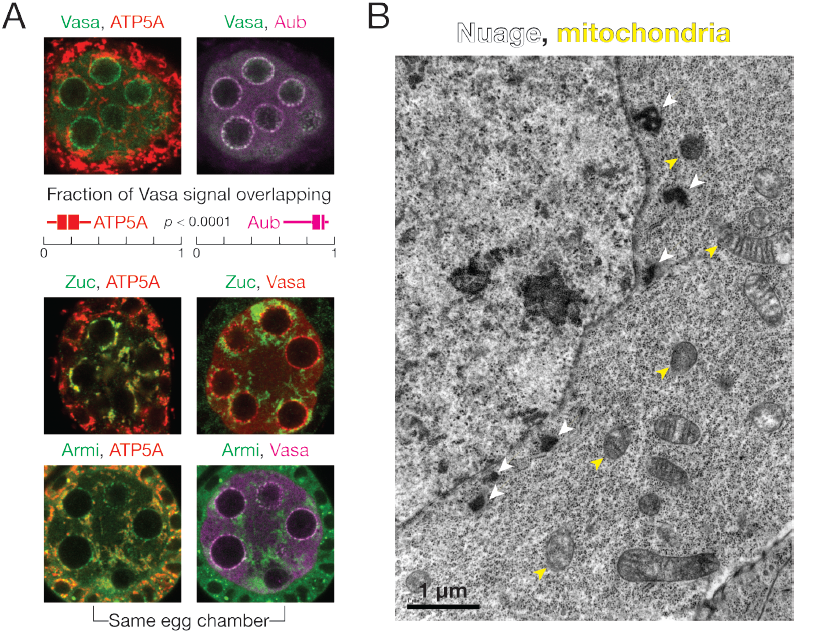
Armi localizes to both nuage and mitochondria, physically separate sites of piRNA biogenesis. (A) Immunofluorescence detection of Vasa, ATP5A and Aub in wild-type stage 3 egg chambers. Box plots show the fraction of Vasa signal overlapping with ATP5A or Aub. Box comprises the 25^th^ percentile and the 75^th^ percentile of data, with the median indicated; whiskers mark 1.5 × interquartile range. p-value was calculated using the twotailed Wilcoxon matched-pairs signed rank test. Immunofluorescence of Zuc-3×FLAG or Armi to detect their colocalization with ATP5A or Vasa in wild-type stage 3 egg chambers. (B) Transmission electron microscopy image of a wild-type stage 3 egg chamber.

### Zuc Localizes to Mitochondria, but not Nuage

In cultured ovarian somatic cells, tagged, overexpressed Zuc localizes to mitochondria through its N-terminal mitochondrial localization signal peptide (Saito et al., 2010; Handler et al., 2013). We examined the localization of endogenous Zuc in the germline using a fly strain in which endogenous Zuc bears a C-terminal 3×FLAG tag (Ge et al.,2016). Homozygous flies expressing only Zuc-3×FLAG showed no detectable defects in fertility or piRNA biogenesis, and the tagged Zuc, expressed from its native chromosomal location, was readily detected using anti-FLAG antibody. Essentially all anti-FLAG immunofluorescence coincided with ATP5A; little anti-FLAG staining overlapped with Vasa (Figure 1A, middle row). We conclude that within the limits of immunofluorescent detection, Zuc localizes to mitochondria and not nuage.

These experiments cannot exclude the possibility that a small but biologically important population of Zuc is present in nuage and contributes to piRNA biogenesis. To test this possibility, we asked whether Zuc interacts with nuage proteins. We immunoprecipitated 3×FLAG-tagged endogenous Zuc from ovary lysates and identified co-immunoprecipitating proteins by mass spectrometry. Under native conditions, mass spectrometry failed to detect any known piRNA biogenesis proteins co-immunoprecipitating with Zuc. Because Zuc-interacting proteins may dissociate upon cell lysis, we surveyed a series of membrane-permeable, reversible chemical crosslinkers for their ability to stabilize Zuc protein-protein interactions. Crosslinking has the added benefit of withstanding stringent washes, thus ensuring that any detected interactions occurred in the cell and not after lysis (Mili and Steitz, 2004). The mouse Armi homolog Mov10l1 co-immunoprecipitates with PLD6, the mouse Zuc homolog, when the two proteins are co-expressed in cultured mammalian somatic cells (Vourekas et al., 2015). Therefore, we tested whether Armi co-immunoprecipitated with Zuc using ovaries treated before lysis with the primary amine-to-primary amine crosslinkers paraformaldehyde, disuccinimidyl tartrate, ethylene glycol bis(succinimidyl succinate), or dithiobis(succinimidyl propionate), or the sulfhydryl-to-sulfhydryl crosslinker dithio-bis-maleimidoethane (DTME). Among these, DTME best stabilized the interaction between Zuc and Armi (Figure S1C).

To more comprehensively identify Zuc-associated proteins, 3×FLAG-tagged Zuc was immunoprecipitated from DTME-crosslinked ovaries, and Zuc-associated proteins identified by mass spectrometry (Table 1). Among the 39 proteins reproducibly co-immunoprecipitated with Zuc (*n* = 3), 17 were mitochondrial proteins not known to participate in piRNA biogenesis. An additional four were piRNA pathway proteins known to localize to mitochondria, including the phased piRNA biogenesis proteins Armi, Gasz, and Minotaur (Olivieri et al., 2010; Saito et al., 2010; Baena-Lopez et al., 2013; Handler et al., 2013; Vagin et al., 2013), as well as the mitochondrial outer membrane protein Papi, which is required both to maintain piRNA levels and to trim pre-piRNAs (Honda et al., 2013; Han et al., 2015a; Hayashi et al., 2016; Izumi et al., 2016; Nishida et al.,2018). Sister of Yb (SoYb), a piRNA pathway protein that colocalizes with Armi to cytoplasmic clouds likely to be mitochondria (Handler et al., 2011), also co-purified with Zuc. In all, 21 proteins known to localize to mitochondria or function in phased piRNA production co-purified with Zuc, strong evidence that phased piRNAs are produced on the outer surface of mitochondria. In contrast, only a single known component of nuage was enriched in the Zuc immunoprecipitate: Armi.

**Table 1.** All data are mean ± SD for weighted spectra count (*n* = 3) and FDR < 0.05 for experimental immunoprecipitate (IP) versus control. (**A**) piRNA biogenesis proteins specifically co-immunoprecipitated with Zuc, Armi or Armi^K729A^. (**B**) Proteins not known to function in piRNA biogenesis that co-immunoprecipitated with Zuc.

| Protein | Localization | Zuc IP | Control IP for Zuc | Armi IP | Armi<sup>K729A</sup> IP | Control IP for Armi |
| --- | --- | --- | --- | --- | --- | --- |
| Zucchini (Zuc) | Mitochondria | 31 ± 6 | 0 | – | – | – |
| Gasz | Mitochondria | 15 ± 5 | 0 | 7 ± 2 | 8.3 ± 0.6 | 0 |
| Papi | Mitochondria | 11 ± 1 | 0 | – | – | – |
| Armitage (Armi) | Mitochondria & nuage | 38 ± 3 | 5 ± 4 | 200 ± 90 | 180 ± 80 | 10 ± 20 |
| Minotaur (Mino) | Mitochondria & ER | 16 ± 4 | 0.7 ± 0.6 | 11 ± 5 | 14 ± 2 | 0.3 ± 0.6 |
| Sister of Yb (SoYb) | Mitochondria? | 4 ± 4 | 0 | 10 ± 10 | 20 ± 10 | 0 |
| Vreteno (Vret) | Nuage | – | – | 4 ± 5 | 4 ± 5 | 0 |
| Shutdown (Shu) | Nuage | – | – | 8 ± 4 | 16 ± 5 | 1 ± 1 |
| Argonaute 3 (Ago3) | Nuage | – | – | 7 ± 7 | 4 ± 5 | 0.3 ± 0.6 |
| Spindle E (Spn-E) | Nuage | – | – | 9 ± 9 | 9 ± 9 | 1 ± 2 |
| Tapas | Nuage | – | – | 8 ± 4 | 8 ± 2 | 2 ± 2 |
| Aubergine (Aub) | Nuage | – | – | 40 ± 20 | 40 ± 20 | 10 ± 7 |
| Brother of Yb (BoYb) | Nuage | – | – | 5 ± 6 | 3 ± 4 | 0.3 ± 0.6 |
| Nibbler (Nbr) | Nuage | – | – | 10 ± 4 | 14 ± 3 | 3.3 ± 0.6 |
| Qin | Nuage | – | – | 5 ± 7 | 2 ± 4 | 0 |
| P-element induced wimpy testis (Piwi) | Nucleus | – | – | 16 ± 6 | 20 ± 10 | 9 ± 6 |
| Vasa (Vas) | Nuage | – | – | 26 ± 6 | 33 ± 2 | 14 ± 2 |
| Tudor (Tud) | Nuage | – | – | 40 ± 30 | 30 ± 30 | 20 ± 20 |

(B) Proteins not known to function in piRNA biogenesis that co-immunoprecipitated with Zuc.
| Mitochondrial Proteins |  |  |  | Others |  |  |  |
| --- | --- | --- | --- | --- | --- | --- | --- |
| Fly name | Human name | Zuc IP | Control IP | Fly name | Human name | Zuc IP | Control IP |
| Miro | Rhot1 | 10 ± 3 | 0 | CG12360 | Trabd | 12 ± 5 | 0 |
| Miga | Miga2 | 11 ± 2 | 1 ± 1 | CG10880 | – | 11 ± 3 | 0.3 ± 0.6 |
| CG9855 | March5 | 5 ± 2 | 0 | nmd | Atad1 | 16 ± 6 | 1 ± 1 |
| Spoon/Yu | Akap1 | 10 ± 4 | 0.7 ± 0.6 | Aldh-III | Aldh3b1 | 9 ± 2 | 0 |
| Marf | Mfn2 | 27 ± 4 | 2 ± 2 | CG7705 | – | 10 ± 3 | 0 |
| Usp30 | Usp30 | 3 ± 4 | 0 | Faf | Faf2 | 7 ± 5 | 0 |
| Men | Me1 | 13 ± 9 | 1 ± 2 | Pex3 | Pex3 | 5 ± 1 | 0.3 ± 0.6 |
| Mtch | Mtch2 | 12 ± 6 | 2 ± 2 | CG2316 | Abcd2 | 5 ± 2 | 0 |
| Ptp61F | Ptpn1 | 5 ± 1 | 0.3 ± 0.6 | Myt1 | PKMYT1 | 4 ± 2 | 0 |
| CG7639 | SAMM50 | 4 ± 3 | 0 | CG1291 | Alg2 | 5 ± 3 | 0.3 ± 0.6 |
| CG1665 | Marc1 | 15 ± 7 | 1.3 ± 0.6 | CG8735 | LNP | 6.3 ± 0.6 | 0.3 ± 0.6 |
| CG7461 | Acadvl | 3 ± 1 | 0 | CG6550 | CDKAL1 | 4 ± 3 | 0 |
| Tom70 | Tomm70 | 28 ± 7 | 5 ± 1 | Sec63 | Sec63 | 7 ± 3 | 1 ± 2 |
| CG3394 | SLC27A1 | 5 ± 4 | 1 ± 1 | CG9853 | Get4 | 5 ± 3 | 0.3 ± 0.6 |
| Pmp70 | Abcd3 | 11 ± 5 | 2 ± 3 | CG6744 | Exd2 | 3 ± 4 | 0 |
| iPLA2-VIA | Pla2g6 | 7 ± 3 | 1.3 ± 0.6 | CG7546 | Bag6 | 10 ± 6 | 2 ± 1 |
| Acsl | Acsl3 | 17 ± 2 | 7 ± 2 | CG6089 | Tmem214 | 13 ± 6 | 2 ± 1 |

### Armi-Associated Proteins Suggest Armi Shuttles from Nuage to Mitochondria

For pre-pre-piRNA to be made in the nuage and processed into phased piRNAs on mitochondria, such piRNA precursors need to be delivered from nuage, across the cytoplasm, to the outer face of mitochondria. A likely candidate to serve this role is Armi, a protein found in both compartments (Cook et al., 2004; Handler et al., 2013; Huang et al., 2014; Pandey et al., 2017). To test whether Armi participates in piRNA biogenesis in both nuage and mitochondria, ovaries from flies producing transgenic 3×FLAG-(Myc)6- tagged Armi (FLAG-MYC-Armi) in the germline were crosslinked with DTME before cell lysis, followed by FLAG immunoprecipitation and mass spectrometry. Flies of the same genetic background, but not carrying the transgene served as the control. Of the 95 proteins reproducibly associated with Armi (*n* = 3), just two (Clueless and Mitochondrial assembly regulatory factor) were mitochondrial proteins without functions in piRNA biogenesis (Table S1). In contrast, 17 were known piRNA factors (Table 1): the mitochondrial proteins Gasz, Minotaur, SoYb, and Papi; the nuage proteins Vreteno, Shutdown, Ago3, Spindle E (Spn-E), Tapas, Aub, Brother of Yb, Nibbler, Qin, Vasa, and Tudor; and the trailing piRNA-guided protein Piwi. Piwi has been previously shown to interact with Armi (Haase et al., 2010; Olivieri et al., 2010; Saito et al., 2010). We conclude that Armi participates in the piRNA pathway both in nuage and on mitochondria.

In theory, Armi might localize to different subcellular compartments at different stages of egg chamber maturation, or Armi may localize to both nuage and mitochondria within a single nurse cell. To differentiate between these two possibilities, we examined the intracellular location of Armi, ATP5A and Vasa in individual stage 3 egg chambers. Armi colocalized with both ATP5A and Vasa in every nurse cell examined (*n* = 60; Figure 1A, bottom row). Thus, Armi is likely to participate in the delivery of pre-pre-piRNAs from nuage to mitochondria.

### Loss of Phased piRNA Production Depletes Armi from the Nuage

If Armi delivers pre-pre-piRNAs from nuage to mitochondria, but plays no other role in piRNA amplification, then loss of Armi should block production of phased piRNAs without perturbing ping-pong. Consistent with this prediction, removing Armi genetically or by germline RNAi disrupts piRNA phasing but not ping-pong piRNA amplification (Handler et al., 2011; Han et al., 2015a). Loss of Zuc similarly disrupts piRNA phasing but not ping-pong (Han et al., 2015a).

Recycling of Armi from mitochondria appears to require phased piRNA production. First, compared to wild type, less Armi was associated with nuage and more with mitochondria in *zuc^H169Y^* mutant ovaries, which produce appropriately localized but catalytically inactive Zuc (Figure 2A). Armi was similarly depleted from nuage but not mitochondria in *minotaur* mutant ovaries (Figure 2A). These data suggest that recycling of Armi from mitochondria back to nuage requires phased piRNA production. In support of the idea that wild-type Armi distribution requires both piRNA amplification and phased piRNA synthesis, inactivating the ping-pong pathway by mutating the catalytic triad of Ago3 from DDH to AAH increases the amount of Armi associated with Ago3, likely in the nuage (Huang et al., 2014).

**Figure 2.**
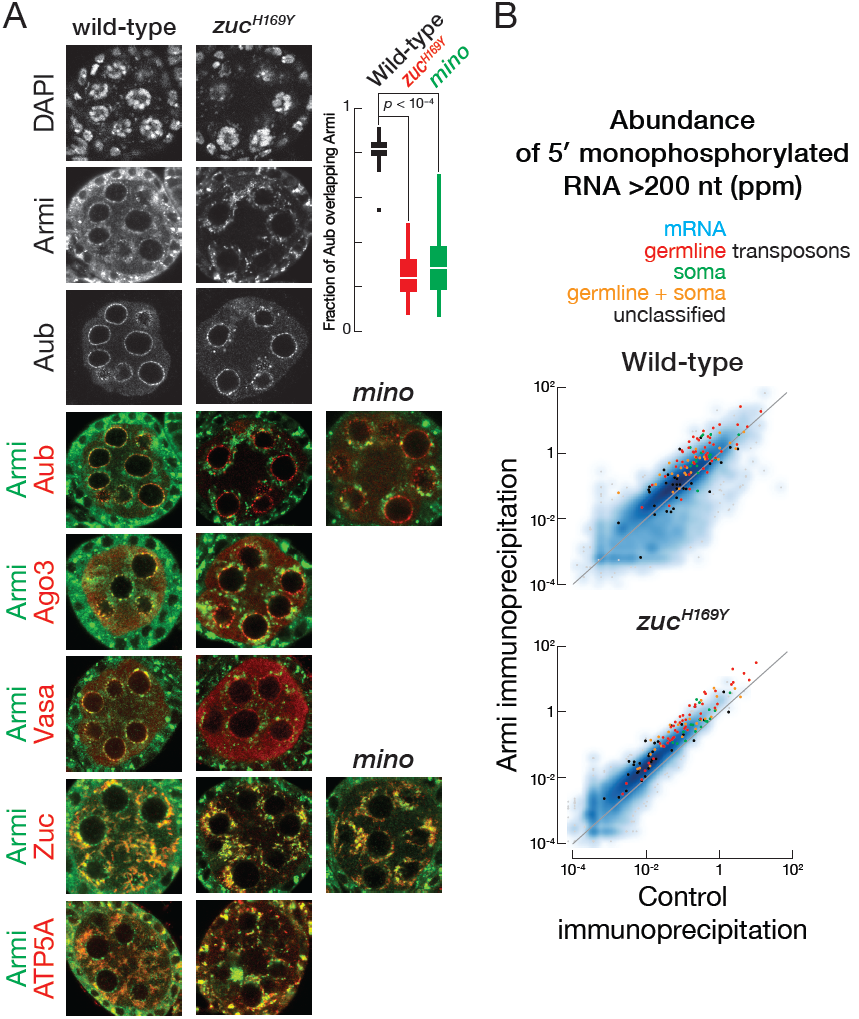
Phased piRNA production promotes nuage localization of Armi. (A) Immunofluorescence co-detection of Armi and Aub, Ago3, Vasa, Zuc-3×FLAG or ATP5A in wild-type or *zuc^H169Y^* stage 3 egg chambers or of Armi and Aub or Zuc-3×FLAG in *minotaur* stage 3 egg chambers. Box plots show the fraction of Aub signal overlapping with Armi in wild-type, *zuc^H169Y^* or *minotaur* stage 3 egg chambers. Box comprises the 25^th^ percentile and the 75^th^ percentile of data, with the median indicated; whiskers mark 1.5 × interquartile range. *p*-value was calculated using the two-tailed Mann Whitney test. (B) Scatter plots of the abundance of transposon- or gene-mapping 5’ monophosphorylated RNA >150 nt co-immunoprecipitated with Armi or control in wild-type or *zuc^H169Y^* mutant ovaries.

### Armi Binds Pre-pre-piRNAs During Ping-pong and Phased piRNA Biogenesis

If Armi delivers pre-pre-piRNAs from nuage to mitochondria, then Armi should bind pre-pre-piRNAs. To test this prediction, we immunoprecipitated FLAG-Myc-Armi from the germline and sequenced associated RNAs >150 nt long and bearing a 5’ monophosphate, the terminal structure found on pre-pre-piRNAs (Han et al., 2015a; Gainetdinov et al., 2018). Ovaries were crosslinked with paraformaldehyde before lysis to ensure detection of only protein-RNA interactions present in vivo. Ovaries without the FLAG-Myc-Armi transgene served as a negative control. Because FLAG-Myc-Armi was expressed only in the germline, sequencing data was normalized to the background of reads mapping uniquely to the somatic follicle cell-specific piRNA cluster *flamenco*. Immunoprecipitated Armi was specifically associated with transposon mapping RNAs with the characteristics of pre-pre-piRNAs (Figure 2B). First, antisense transposon-derived reads were more abundant in the Armi immunoprecipitate (61% of transposon mappers were antisense) than in the control (45% antisense), and, second, the Armi-associated RNA favored a 5’ uridine (Figure 3A).

**Figure 3.**
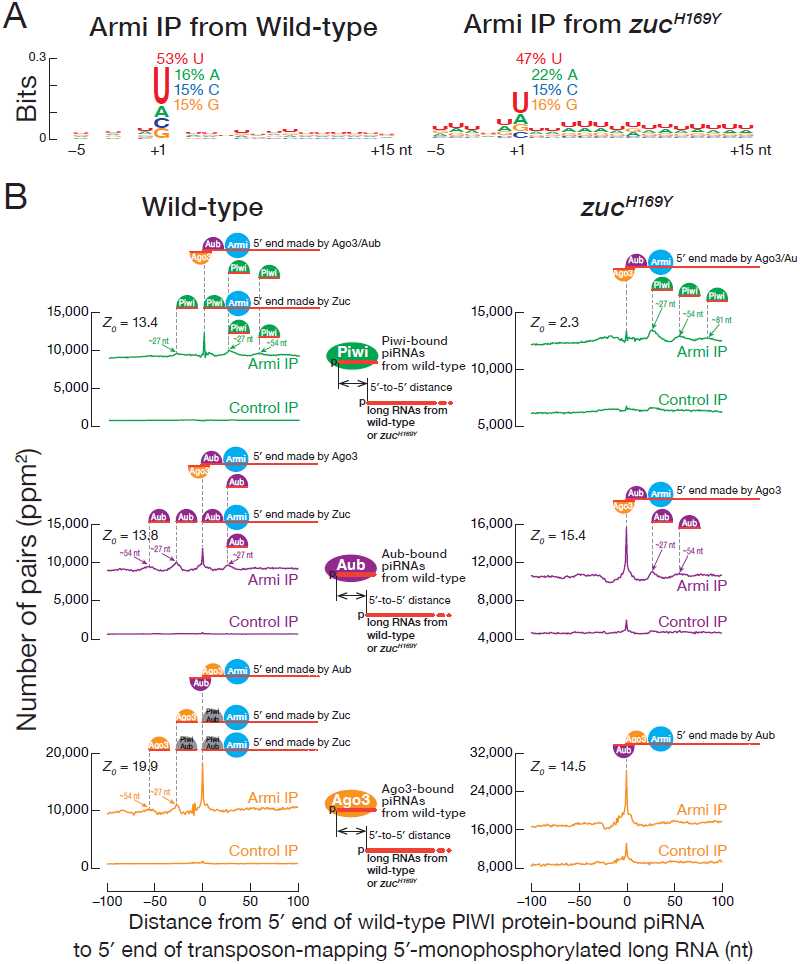
Armi interacts with pre-pre-piRNA during the nuage and mitochondrial phases of piRNA production. (A) Genomic nucleotide bias around the 5’ ends (nt position 1) of 5’ monophosphorylated antisense transposon RNA >150 nt co-immunoprecipitated with Armi. Each RNA 5’ end was weighted equally, ignoring abundance. (B) Distance on the same genomic strand from 5’ ends of PIWI protein-bound piRNAs from wild-type ovaries to 5’ ends of 5’ monophosphorylated long RNAs co-immunoprecipitated with Armi or control in wild-type or *zuc^H169Y^* mutant ovaries.

The current model for *Drosophila* germline piRNA biogenesis posits that in the nuage, an initiator piRNA directs Ago3 (and to a lesser extent, Aub) to cleave a piRNA precursor transcript, generating a 5’ monophosphorylated end that binds Aub. This Aub-bound pre-pre-piRNA is then transferred to the mitochondrial surface to be cleaved by Zuc. Zuc cleavage generates an Aub-bound responder pre-piRNA from the 5’ end of the pre-pre-piRNA, while the rest of the pre-pre-piRNA is processed into a chain of trailing piRNAs loaded into Piwi and, to a smaller extent, Aub. The hypothesis that Armi transports the Aub-bound pre-pre-piRNA from nuage to mitochondria predicts that Armi will interact with two types of transposon-derived 5’ monophosphorylated RNA: (1)RNAs whose 5’ ends are established by Ago3 or Aub cleavage, i.e., ping-pong piRNA precursors; and (2) RNAs whose 5’ ends are made by Zuc and that generate Piwi- or Aub-bound trailing piRNAs.

Our data suggest that Armi binds pre-pre-piRNAs produced both in the ping-pong and phased piRNA biogenesis pathways. First, we measured the distance between the 5’ ends of Piwi-bound piRNAs and the 5’ ends of transposon-mapping, 5’ monophosphorylated RNAs—i.e., putative pre-pre-piRNAs—co-immunoprecipitated with Armi. The 5’ ends of RNAs associated with Armi often coincided with a Piwi-bound piRNA (*Z_0_* = 13.4; Figure 3B, left). That is, Piwi-bound, phased piRNAs typically arise from the 5’ ends of Armi-bound pre-pre-piRNAs (Figure 3B, left). Second, the 5’ ends of RNAs co-immunoprecipitated with Armi also frequently coincided with an Aub- or Ago3-bound piRNA (*Z_0_* = 13.8 or 19.9, respectively; Figure 3B, left), indicating that Armi associates with the immediate precursors of responder piRNAs.

The current model also predicts that in the absence of Zuc, only pre-pre-piRNAs arising from ping-pong slicing are produced. That is, loss of Zuc should restrict pre-pre-piRNA generation to the precursors whose 5’ ends are generated by Ago3 or Aub cleavage. Compared to wild-type Armi-associated long RNA, fewer 5’ ends of the Armi-associated RNAs from *zuc^H169Y^* ovaries coincided with Piwi-bound piRNAs from wild-type, consistent with loss of pre-pre-piRNAs whose 5’ ends are produced by Zuc (Figure 3B, right). Instead, the 5’ ends of the Armi-associated RNAs from *zuc^H169Y^* most often coincided with Aub- or Ago3-bound piRNAs from wild-type, indicating that pre-pre-piRNAs made by ping-pong are bound to Armi in *zuc^H169Y^* ovaries (Figure 3B, right). Because trailing piRNAs are not produced in *zuc^H169Y^* mutants, the 5’ ends of the Piwi- or Aub-bound, but not Ago3-bound piRNAs from wild-type ovaries map 27 nt or 54 nt (one or two pre-piRNA lengths) downstream from the 5’ ends of the pre-pre-piRNAs associated with Armi in *zuc^H169Y^* mutant (Figure 3B, right). In fact, pre-pre-piRNAs whose 5’ ends correspond to an Aub- and Ago3-bound responder piRNAs accumulate in *zuc^H169Y^* (∼50% and ∼80% increase, respectively, compared to wild-type). That is, the pre-pre-piRNAs whose 5’ ends correspond to Piwi- or Aub-bound trailing piRNAs are not generated without Zuc function. We conclude that in addition to binding pre-pre-piRNAs that fuel phased piRNA production, Armi also binds pre-pre-piRNAs typically generated by Ago3 or Aub directed by an initiator piRNA-the piRNA pathway intermediates that must be delivered from the nuage to the mitochondria.

### Armi ATPase Activity Enables Correct substrate Selection

The Armi helicase domain belongs to the Upf1-like superfamily 1. Modeling the Armi helicase core predicts that its structure is highly similar to human Upf1 (RMSD = 0.91 Å, Figure 4A). Upf1 is a core factor in nonsense-mediated mRNA decay (NMD), and mutating lysine 498 to alanine in helicase motif I of human Upf1 disrupts ATP- but not RNA-binding; Upf1^K498A^ loses its selectivity for mRNAs with premature stop codons and instead binds RNA promiscuously (Weng et al., 1996; Weng et al., 1998; Lee et al., 2015).

**Figure 4.**
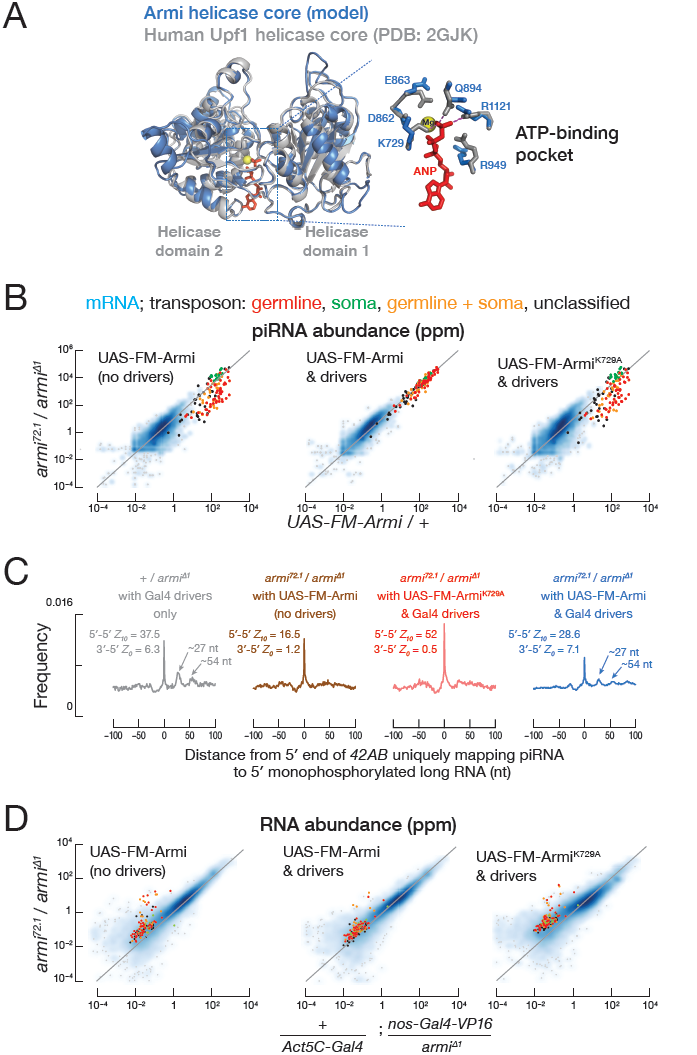
Armi^K729A^ ATPase mutant fails to support phased piRNA production. (A) Predicted Armi helicase core superimposed on the human Upf1 helicase core (Cheng et al., 2007). The enlarged view of the Armi ATP-binding pocket shows amino acids surrounding adenosine-5’-(β,γ-imido)triphosphate and a magnesium ion present in the Upf1 structure. (B) Scatter plot of the abundance of transposon- or gene-mapping piRNAs from the indicated genotype. (C) Distance on the same genomic strand from 5’ ends of piRNAs uniquely mapping to 42AB to the 5’ ends of 5’ monophosphorylated RNA >150 nt from the indicated genotype. *Z*-scores indicate the significance of the 5’-to-5’ distance = 10 nt on opposite genomic strands (ping-pong, *Z_10_*) and the 3’-to-5’ distance = 0 nt on the same genomic strand (phasing, *Z_0_*) for piRNAs used in the analyses. (D) Scatter plot of the abundance of transposon- or gene-mapping RNA >150 nt or piRNAs from the indicated genotype.

The corresponding amino acid substitution in Armi, K729A, does not support piRNA biogenesis: unlike the transgene expressing wild-type Armi, transgenic Armi^K729A^ failed to rescue piRNA production in *armi^72.1/Δ1^* ovaries, which lack germline Armi. Compared to the wild-type transgene, the abundance of piRNAs from germline-specific transposons fell nearly tenfold in Armi^K729A^ ovaries (median = 11%, *n* = 47), a level statistically indistinguishable from *armi^72.1/Δ1^* alone (median = 9%; Wilcoxon matched-pairs signed rank test *p* = 0.07; Figure 4B). Like *armi^72.1/Δ1^*, Armi^K729A^ ovaries produced mainly ping-pong piRNAs (Figure 4C). In contrast, *armi^72.1/Δ1^* rescued with a wild-type Armi transgene made both ping-pong and phased piRNAs (Figure 4C). Unlike wild-type transgenic Armi, the Armi^K729A^ transgene failed to support germline transposon silencing (Figure 4D), and the flies showed defects in both egg laying and hatching (Figure S2A). The defect does not reflect a dearth of Armi^K729A^: the abundance of the transgenic wild-type and K729A mutant proteins were comparable (Figure S2B). Neither increasing nor decreasing Armi abundance relative to the endogenous level had a measurable effect on piRNA levels, demonstrating that Armi abundance does not limit piRNA production (Figure S2C).

To test whether Armi^K729A^ interacts with piRNA precursors, we sequenced 5’ monophosphorylated long RNA co-immunoprecipitated with transgenic Armi^K729A^ in the germline of wild-type ovaries. Compared to transgenic wild-type Armi, Armi^K729A^ bound 18-fold more mRNA degradation products, as judged by the Armi^K729A^ median enrichment over wild-type Armi (Figure 5A). Both Armi^K729A^ and wild-type Armi bound germline transposon-derived pre-pre-piRNAs (Figures 2B and Figure 5A). These data suggest that, like Upf1^K498A^, Armi^K729A^ loses substrate selectivity when it cannot bind ATP.

**Figure 5.**
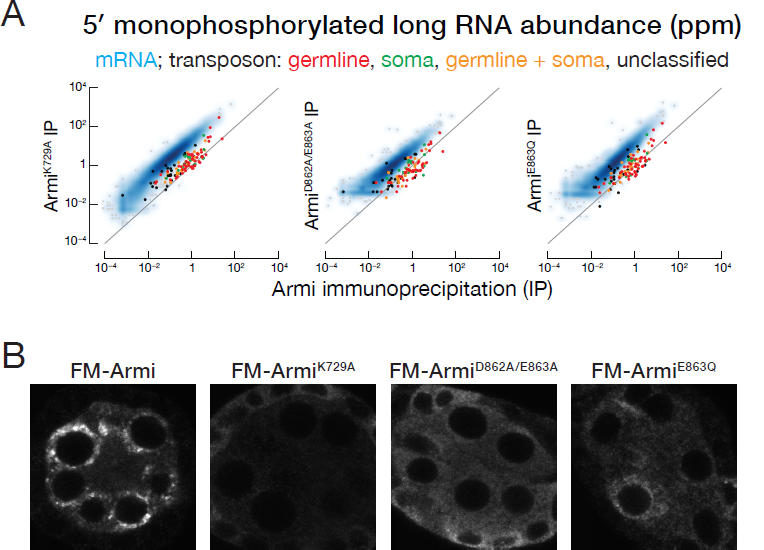
Armi ATPase activity enables correct substrate selection. (A) Scatter plot of the abundance of transposon- or gene-mapping 5’ monophosphorylated RNA >150 nt co-immunoprecipitated with transgenic FLAG-Myc-Armi^K729A^, FLAG-Myc-Armi^DE862863AA^, FLAG-Myc-Armi^E863Q^ versus FLAG-Myc-Armi. (B) Immunofluorescence detection using anti-FLAG antibody of transgenic FLAG-Myc-Armi, FLAG-Myc-Armi^K729A^, FLAG-Myc-Armi^DE862863AA^ or FLAG-Myc-Armi^E863Q^ in *armi* mutant germ cells from stage 3 egg chambers.

Artificially tethering Armi to a transcript triggers piRNA production (Pandey et al., 2017; Rogers et al., 2017). Yet the mRNA degradation products associated with Armi^K729A^ do not fuel an increase in mRNA-derived piRNAs (Figure S2C). Why do the mRNA fragments associated with Armi^K729A^ fail to enter the phased piRNA pathway? Perhaps ATP-binding or hydrolysis by Armi is required for additional steps in phased piRNA biogenesis. Alternatively, mRNA degradation products may be associated with proteins that prevent their delivery to mitochondria.

Another Upf1 mutant, Upf1^D637A/E638A^, still binds ATP but binds RNA promiscuously (Lee et al., 2015). Similarly, mutant Armi bearing the corresponding amino acid substitution, D862A/E863A, inappropriately bound mRNA degradation products (Figure 5A). Finally, mutation of the DEAD motif (E339Q) of the DEAD-box protein Vasa slows release of the products of ATP hydrolysis and traps Vasa on RNA (Xiol et al., 2014). The analogous amino acid substitution in Armi, E863Q, also increased the association of Armi with mRNA degradation products (Figure 5A). We conclude that ATP binding and hydrolysis, as well as subsequent release of the resulting ADP and inorganic phosphate, are required for Armi to discriminate between pre-pre-piRNAs intended to produce transposon-silencing piRNAs and mRNA degradation products that are normally excluded from the piRNA pathway.

Unlike the punctate perinuclear localization of transgenic wild-type Armi, Armi^K729A^ was dispersed throughout the cytoplasm (Figure 5B). Yet Armi^K729A^ remains associated with the same set of piRNA pathway proteins that co-immunoprecipitate with wild-type Armi (Table 1), including the nuage proteins Vreteno, Shutdown, Ago3, Spn-E, Tapas, Aub, Brother of Yb, Nibbler, Qin, and Vasa, and the mitochondrial proteins Gasz, Minotaur, SoYb and Papi. These data suggest that RNA-protein rather than protein-protein interactions with Armi determine its subcellular localization. supporting this view, both Armi^D862A/E863A^ and Armi^E863Q^ (Figure 5B), as well as other mutations in the Armi ATPase catalytic or ATP-binding site (Pandey et al., 2017), cause Armi to be partially dispersed in the cytoplasm. We hypothesize that the loss of pre-pre-piRNA selectivity leads to Armi inappropriately binding mRNA degradation products, in turn causing it to disperse throughout the cytoplasm.

## DISCUSSION

### Armi Moves between Nuage and Mitochondria

Our data suggest that Armi links the nuage and mitochondrial phases of piRNA biogenesis by associating with piRNA precursors and protein factors in both compartments. We propose that pre-pre-piRNAs are generated in the nuage by initiator piRNA-directed cleavage, then accompanied by Armi as they transit to the mitochondria, where Zuc generates both ping-pong and phased pre-piRNAs. We cannot currently determine whether transfer of Armi-bound pre-pre-piRNAs is an active process or simply the consequence of passive diffusion. In either case, the interactions of Armi with mitochondrial proteins likely serve to anchor Armi-associated pre-pre-piRNAs to the mitochondrial surface when it arrives at the organelle. An active piRNA pathway is required for release of Armi from nuage or mitochondria: catalytically inactive Ago3, a mutation that block ping-pong cleavages, traps Armi in nuage (Huang et al., 2014), while *zuc^H169Y^*, which cannot generate phased piRNAs, traps Armi on mitochondria (Figure 2A). Finally, in Armi mutants that promiscuously bind mRNA, Armi disperses throughout the cytoplasm.

We envision that Armi first binds to piRNA precursors in the nuage (Figure 6). Once an Ago3-catalyzed ping-pong cleavage generates a 5’ monophosphorylated pre-pre-piRNA bound to Aub, the Armi-Aub-pre-pre-RNA complex is released into the cytoplasm. When it encounters the surface of a mitochondrion, the complex can be retained by protein-protein interactions such as Armi binding to Gasz (Handler et al., 2013). On the mitochondrial outer membrane, Zuc and other piRNA pathway proteins act to release an Aub-bound responder piRNA and to convert the rest of the pre-pre-piRNA into trailing piRNAs bound to Piwi. In this model, complete conversion of a pre-pre-piRNA into piRNAs releases Armi to the cytoplasm, allowing it to recycle to the nuage. It is tempting to speculate that Armi, free of RNA, has a higher affinity for nuage proteins and that Armi gains a higher affinity for mitochondrial proteins when associated with a pre-pre-piRNA. Currently, there is no direct evidence that Armi changes conformation upon binding RNA, but such a model is consistent with current knowledge of Upf1-like proteins.

**Figure 6.**
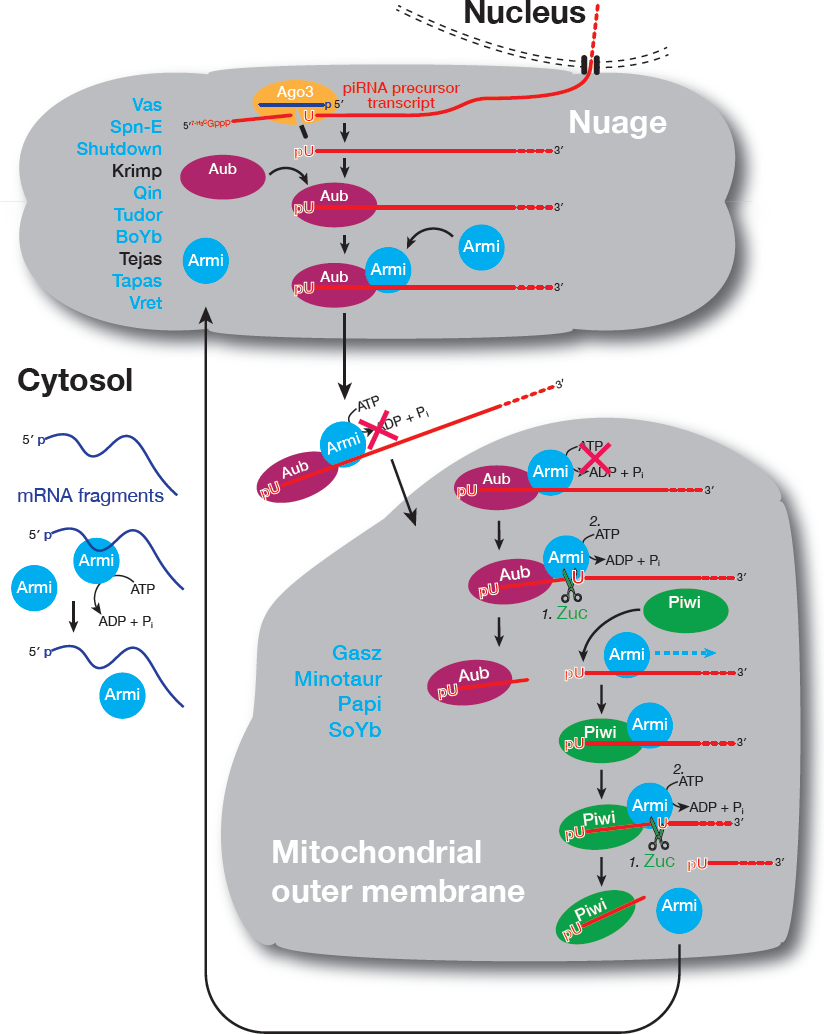
A model for the role of Armi in piRNA biogenesis. In addition to the three PIWI proteins, other piRNA factors interacting with Armi are shown in blue.

The movement of piRNA proteins between nuage and cytoplasm has been reported for Vasa, Tudor, Aub, Tejas, Spn-E and Ago3 (Xiol et al., 2014; Webster et al., 2015; Andress et al., 2016). Except for Tejas, all of these proteins associate with Armi (Table 1A), suggesting that they accompany Armi when it moves from the nuage into the cytoplasm. Interestingly, none of these five proteins co-immunoprecipitated with Zuc, suggesting that they dissociate from the Armi-pre-pre-piRNA complex in the cytoplasm or upon reaching the mitochondrial outer membrane but before Zuc engages Armi (Figure 6). In fly germline nurse cells, nuage and mitochondria are physically separate, a property that facilitated our study of Armi re-localization between the two compartments. However, in some animal germ cells, such as those of mice and frogs, nuage and mitochondria are directly apposed (Eddy, 1974; Eddy, 1975; Kloc et al., 2014). We anticipate that Armi plays a similar role in these species, escorting pre-pre-piRNA generated in nuage to the mitochondria for conversion into piRNAs. However, without the clear separation of nuage and mitochondria of flies, visualizing Armi dynamics will likely be challenging.

### Redundant Roles for Spoon and Papi in piRNA Biogenesis?

At least 127 proteins interact with Zuc or Armi, including 32 uncharacterized proteins (Tables 1 and S1), some of which may correspond to piRNA pathway proteins whose functions overlap with those of known piRNA factors, causing them to be missed by previous screens. For example, Armi interacts with the Tudor domain-containing protein CG9925, which is predicted to be one of the two fly orthologs of mammalian TDRD1 (Table S1). Zuc interacts with the Tudor domain-containing proteins Spoon and Papi, a pair predicted to have similar functions (Handler et al., 2011). Individually, depleting the paralogous proteins CG9925 or CG9684 or the paralogs Spoon or Papi by RNAi does not de-repress transposons, suggesting redundant roles within each pair (Handler et al., 2011). Loss of Papi in cultured *Bombyx mori* (silk moth) BmN4 cells decreases piRNA levels (Izumi et al., 2016; Nishida et al., 2018), but *papi* mutant flies and *Tdrkh* (the mammalian Papi homolog) mutant mice show little change in piRNA abundance (Saxe et al., 2013; Han et al., 2015a). The *B. mori* and mouse genomes both encode Papi paralogs similar to fly Spoon (BGIBMGA006841 and Akap1). The finding that both Spoon and Papi interact with Zuc suggests that Spoon may compensate for loss of Papi in *Drosophila*.

### Armi ATPase Confers substrate Discrimination in piRNA Biogenesis

The promiscuous mRNA binding of Armi ATPase mutants (Figure 5A) calls to mind the aberrant binding of Upf1 ATPase mutants to non-NMD targeted mRNAs (Lee et al., 2015). The Upf1 ATPase activity is proposed to be rapidly activated on non-target mRNAs, promoting dissociation; on NMD targets the ATPase is temporarily inhibited allowing Upf1 to dwell longer (Lee et al., 2015), and ATP or ADP decrease the association between the Upf1 helicase domain and single-stranded RNA (Cheng et al., 2007).

We envision (Figure 6) that in the cytoplasm, the Armi ATPase allows it to rapidly dissociate from genic mRNA; in nuage, Armi ATP-binding or ATPase activity is likely inhibited by interactions with proteins such as Aub, which stabilize Armi binding to pre-pre-piRNAs. After an Armi- and Aub-bound pre-pre-piRNA arrives at the mitochondrial surface, Zuc cleavage liberates an Aub-bound pre-piRNA from the 5’ end of the pre-pre-piRNA, activating the Armi ATPase and allowing Armi to translocate 5’-to-3’ along the pre-pre-RNA. Armi translocation would allow the newly shortened pre-pre-piRNA to bind a new Aub or Piwi at its 5’ end. Binding of Aub or Piwi would then restore inhibition of the Armi ATPase until Zuc makes the next cleavage to liberate a trailing piRNA. Because Aub and Piwi are not freely available in the cytosol, the ATPase activity of delocalized Armi is not inhibited, enabling rapid dissociation of Armi from incorrect RNA substrates, conferring substrate selectivity.

## ACCESSION NUMBERS

Sequencing data are available from the National Center for Biotechnology Information Sequence Read Archive using accession number PRJNA495886.

## ACKNOWLEDGEMENTS

We thank John Leszyk and Michelle Dubuke at the UMass Proteomics Core, Keith Reddig at the UMass Electron Microscopy Core, Christina Baer at the Sanderson Center for Optical Experimentation, and Haiwei Mou, Cha San Koh, and members of the Zamore laboratory for help and discussions. This work was supported by NIH grant R37GM062862 to P.D.Z. and P01HD078253 to Z.W. and P.D.Z. The UMass Electron Microscopy Core Facility is supported by grant S10RR027897 from the National Center for Research Resources.

## AUTHOR CONTRIBUTIONS

D.T.G. and P.D.Z. conceived the experiments. D.T.G., W.W., Z.W. and P.D.Z. designed the experiments. D.T.G. and C.T. performed the experiments. W.W., I.G. and D.T.G. analyzed the sequencing data. D.T.G, Z.W., and P.D.Z. wrote the manuscript.

## SUPPLEMENTAL ITEM TITLES

**Figure S1.**
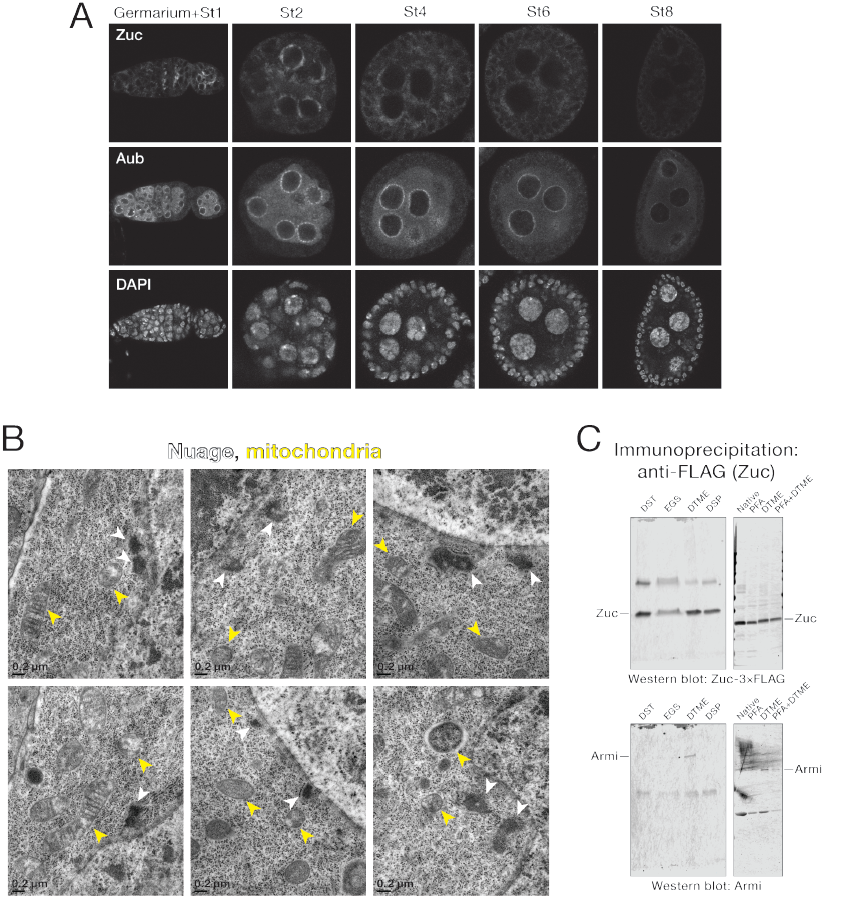
piRNA biogenesis factors are expressed higher in early stages of oogenesis; nuage and mitochondria are separate in nurse cells; DTME best crosslinks Armi to Zuc. Related to Figure 1. (A) Immunofluorescence detection of Zuc-3×FLAG, Aub and nucleic acids (DAPI). St, stage. (B) Transmission electron microscopy of wild-type stage 3 egg chambers. (C) Before tissue lysis, *Zuc-3×FLAG* fly ovaries were crosslinked as indicated. Eluate from the FLAG immunoprecipitation was analyzed by Western blotting.

**Figure S2.**
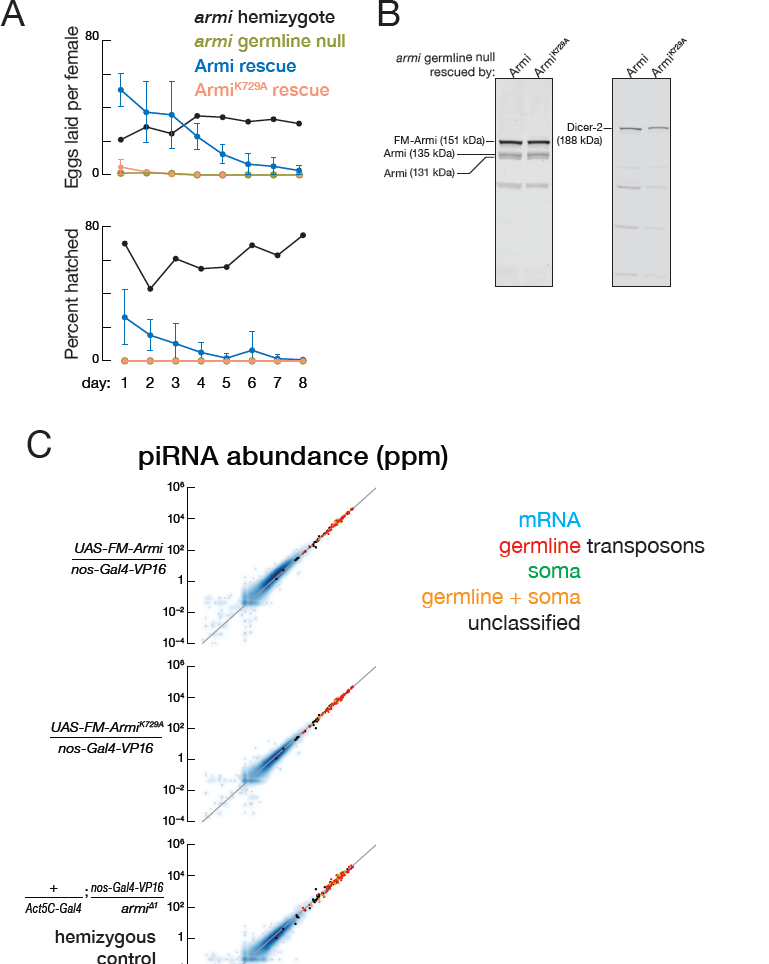
piRNA level in UAS-FM-Armi overexpression, UAS-FM-Armi^K729A^ overexpression, and *armi* hemizygote flies; Armi^K729A^ is expressed at a similar level to Armi. Related to Figure 4. (A) Female fertility test. For UAS-FM-Armi and UAS-FM-Armi^K729A^ rescues, error bars show S.D. from three biological replicates. (B) Western blotting to detect Armi. Dicer-2 provided a loading control. (C) Scatter plot of the abundance of transposon- or gene-mapping piRNAs from the indicated genotype.

**Table S1.**
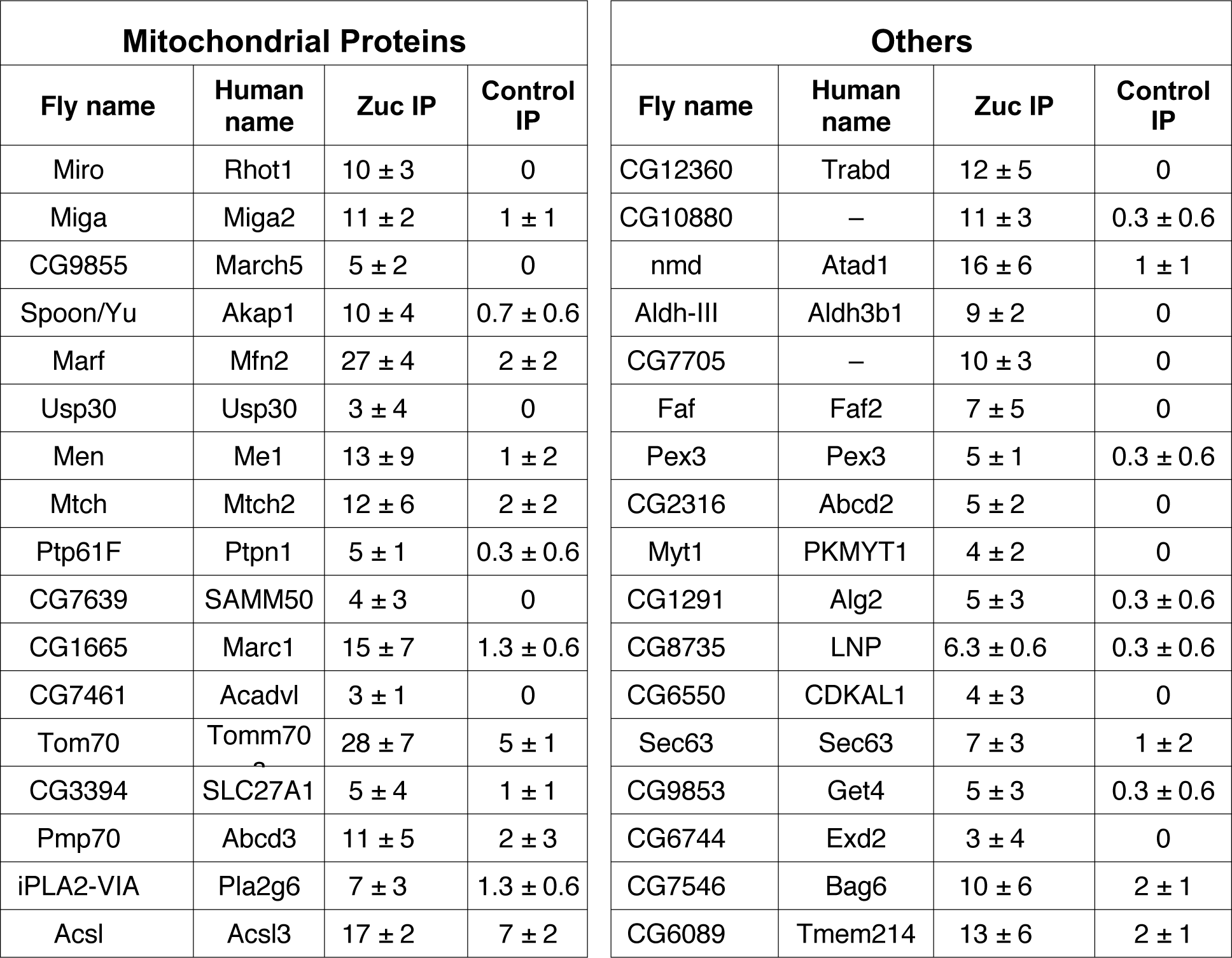
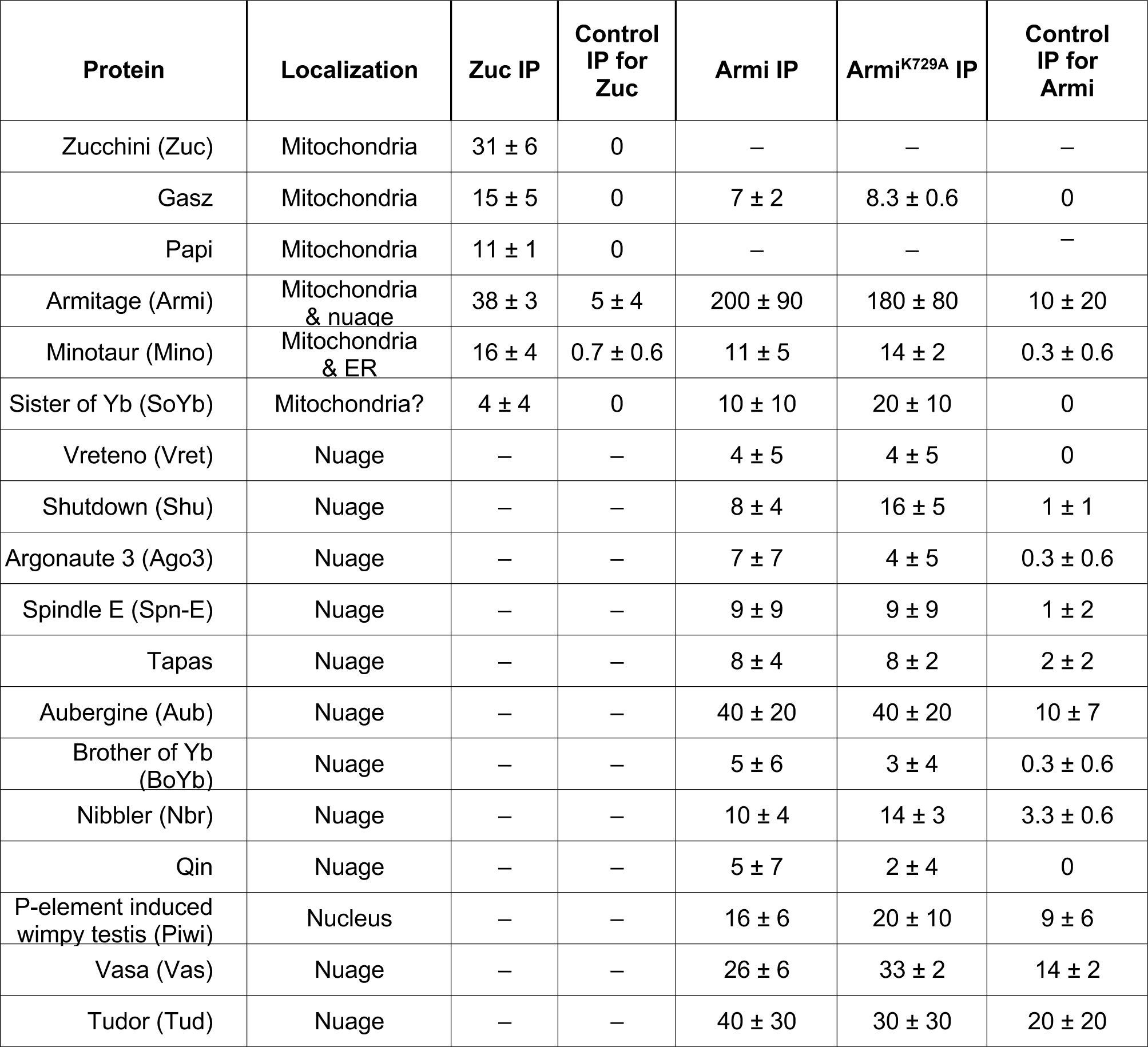
Proteins specifically co-immunoprecipitated with Armi in DTME-crosslinked ovaries (FDR < 0.05). Related to Figure 1. (**A**) piRNA biogenesis proteins specifically co-immunoprecipitated with Zuc, Armi or Armi^K729A^ (**B**) Proteins not known to function in piRNA biogenesis that co-immunoprecipitated with Zuc.

## KEY RESOURCES TABLE

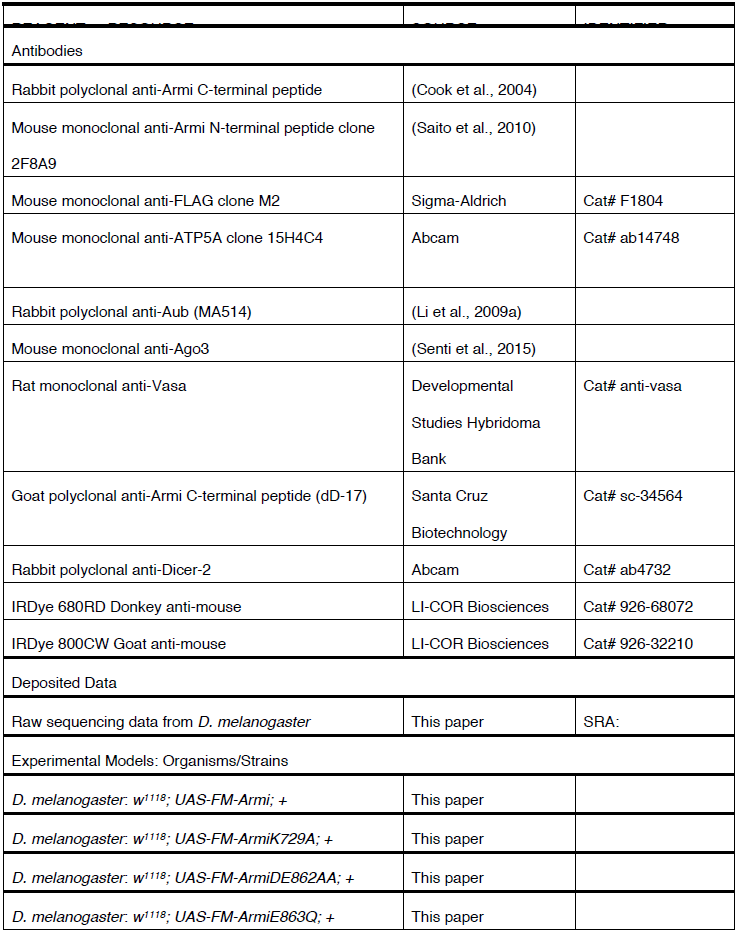
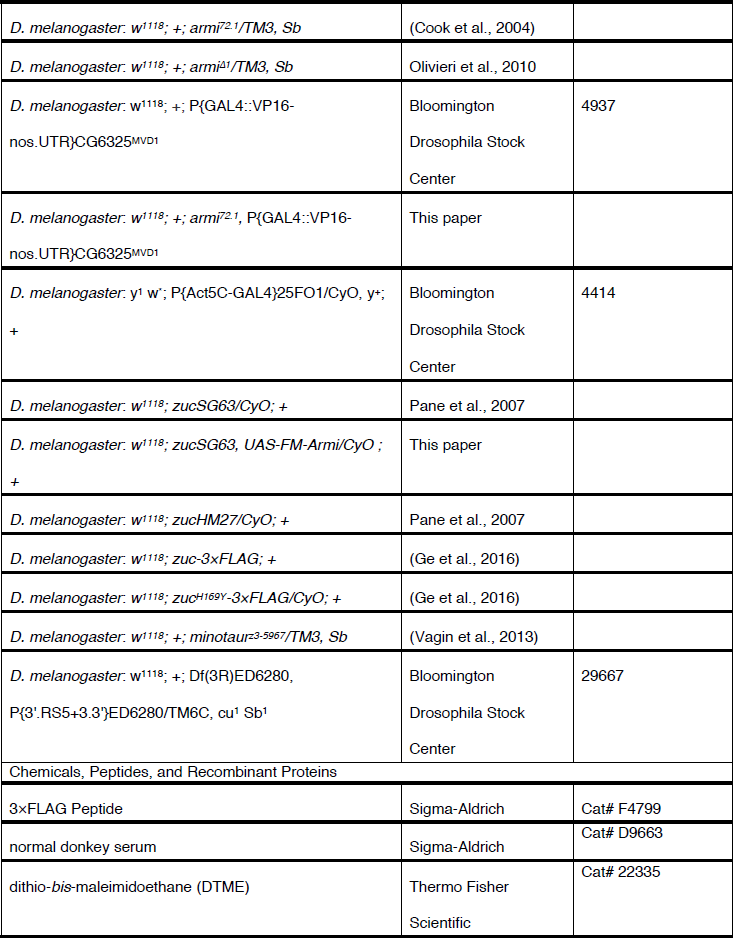
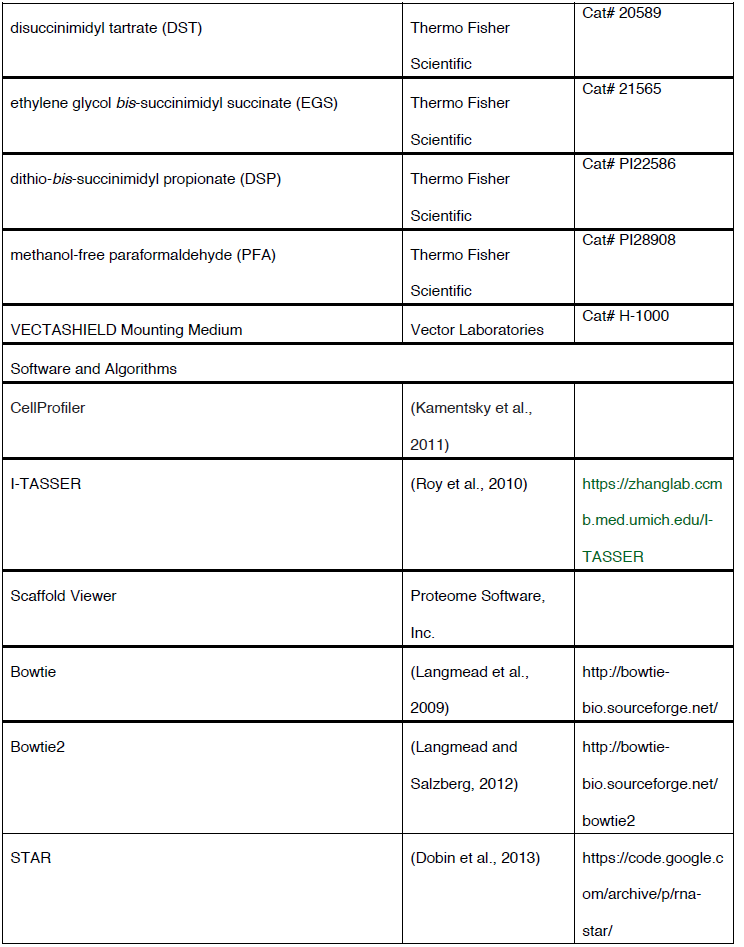
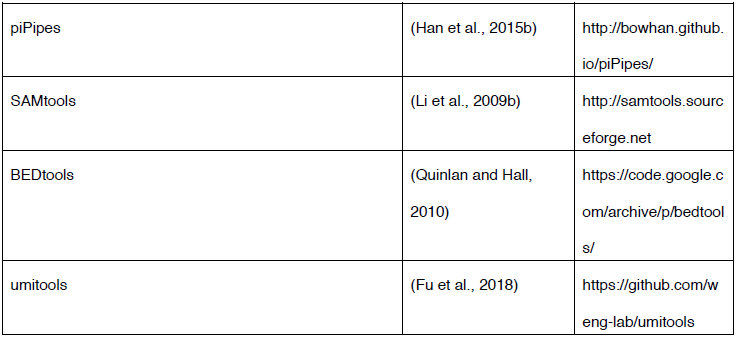

## CONTACT FOR REAGENT AND RESOURCE SHARING

Further information and requests for resources and reagents should be directed to, and will be fulfilled by, the Lead Contact, Phillip D. Zamore, or by completing the request form at www.zamorelab.umassmed.edu/reagents.

## STAR Methods

### Drosophila Stocks

***Transgenic 3×FLAG-6xMyc-Armi:*** cDNA corresponding to *armi* mRNA isoform A (NM_001014556; 3558 bp) was cloned into pENTR-D-TOPO (Invitrogen); K729A, DE862AA and E863Q mutations were generated by site-directed mutagenesis. Wild-type and mutant sequences were subcloned into the modified Gateway vector pPFM-attB, which includes *UASp* sites, a *3×FLAG-6×Myc* N-terminal tag, and an *attB* site for site-specific integration by the *PhiC31* integrase-mediated transgenesis system. The pPFM-attB-armi plasmid was injected into *attP40* flies carrying the *attP* landing site at cytological band 25C7 in chromosome 2L. Rainbow Transgenic Flies, Inc. (Camarillo, CA, USA) performed injections and established transgenic lines *UAS-FM-Armi, UAS-FM-ArmiK729A, UAS-FM-ArmiDE862AA* or *UAS-FM-ArmiE863Q*.

***Rescue of* armi *germline null mutants:*** Armi function is needed in somatic follicle cells for ovary development, and *armi* null flies *(armi^Δ1^)* develop rudimentary ovaries (Olivieri et al., 2010). *armi^72.1^*, an incomplete excision of a *P-element* inserted into the *armi* 5’-untranslated region (UTR), produces somatic but not germline Armi (Olivieri et al., 2010). *armi^Δ1^* removes the *armi, CycJ*, and *CG14971* coding sequences. We therefore used trans-heterozygous *armi^72.1^/armi^Δ1^* as the *armi* germline null mutant background. To test whether the *UAS-FM-Armi* transgene rescues *armi^721^/armi^Δ1^, armi^72.1^* was recombined with the third-chromosome, germline-specific Gal4 driver *P{GAL4::VP16-nos.UTR}CG6325[MVD1]* (Bloomington #4937) to yield *armi^72.1^, nos-Gal4-VP16*. A second-chromosome Gal4 driver, *P{Act5C-GAL4}25FO1* (Bloomington #4414), which is expressed throughout the ovary, was also included to increase *UAS-FM-Armi* expression. Rescue flies had the genotype: *w^1118^; UAS-FM-Armi/Act5C-Gal4; armi^72.1^, nos-Gal4-VP16/armi^Δ1^* (Armi rescue) or *w^1118^; UAS-FM-ArmiK729A/Act5C-Gal4; armi^72.1^, nos-Gal4-VP16/armi^Δ1^* (ArmiK729A rescue).

***Germline-specific overexpression of transgenic Armi: UAS-FM-Armi*** was overexpressed using the third-chromosome, germline-specific Gal4 driver *P{GAL4::VP16-nos.UTR}CG6325[MVD1]*. To overexpress *UAS-FM-Armi* in *zuc^H169Y^* background, *UAS-FM-Armi* was recombined with *zuc^SG63^*, a point mutant allele changing histidine 169 to tyrosine (henceforth *zuc^H169Y^)*, then crossed in trans to *zuc^HM27^*, a null allele (Pane et al., 2007). The same third-chromosome germline-specific Gal4 driver *P{GAL4::VP16-nos.UTR}CG6325[MVD1]* was used to express *UAS-FM-Armi* in a *zuc^H169Y^* background.

***Other stocks:*** Wild-type and the H169Y mutant endogenous Zuc, tagged with 3×FLAG strains have been described previously (Ge et al., 2016). *minotaur^z3-5967^* (Vagin et al., 2013) was crossed in trans to *Df(3R)ED6280* (Bloomington #29667) to obtain *minotaur* mutants.

### Female Fertility

Female fertility was tested as described in (Li et al., 2009a) except five two day-old female virgins were mated at 25°C to three Oregon R males in a small cage (70 mm tall) on a 60 mm diameter grape juice agar plate dabbed with yeast paste (Lesaffre Red Star bakers active dry yeast). The plate was removed from the cage after 24 h, and the number of eggs laid counted. The plate was further incubated at 25°C for 24 h, then the number of eggs hatched were counted. Plates were replaced and scored daily for 8 days.

### Ovary Isolation and Crosslinking

Flies were grown at 25°C. Unless otherwise noted, 0–3 day-old female flies were fed yeast paste for two days before ovary dissection. Ovaries (30–50 mg) were dissected into dissection buffer (5 mM HEPES, pH 7.2, 128 mM NaCl, 2 mM KCl, 4 mM MgCl_2_, 1.8 mM CaCl_2_, 35.5 mM sucrose) and transferred to 1.7 ml microfuge tubes on ice.

Next, dissection buffer was replaced with 1 ml 2 mM dithio-bis-maleimidoethane (DTME, Thermo Fisher #22335), 5 mM disuccinimidyl tartrate (DST, Thermo Fisher #20589), 5 mM ethylene glycol bis-succinimidyl succinate (EGS, Thermo Fisher #21565) or 5 mM dithio-bis-succinimidyl propionate (DSP, Thermo Fisher #PI22586) in dissection buffer or 0.2% (w/v) methanol-free paraformaldehyde (PFA, Thermo Fisher #PI28908) in 0.1 M sodium phosphate, pH 7.3. Crosslinking was performed for 10 min (PFA), 15 min (DTME) or 30 min (DST, EGS, DSP), then the crosslinking solution removed and ovaries washed three times for 5 min with 50 mM Tris-HCl, 150 mM NaCl, pH 7.5 (1× TBS) at room temperature. The ovaries were frozen in liquid nitrogen and stored at −80°C. PFA crosslinking was reversed for protein analysis by heating at 95°C for 30 min for RNA experiments by heating at 65°C for 2 h. DTME crosslinking was reversed by heating at 37°C for 30 min in the presence of 10 mM dithiothreitol (DTT); DST crosslinking was reversed by incubating at room temperature for 30 min in the presence of 15 mM sodium periodate; EGS crosslinking was reversed by heating at 37°C for 3 h in the presence of 1 M hydroxylamine-HCl; DSP crosslinking was reversed by heating at 95°C for 5 min in the presence of 100 mM DTT.

### Immunofluorescence

Intact ovaries were fixed in 4% methanol-free paraformaldehyde (w/v) in 0.1 M sodium phosphate, pH 7.3, for 10 min, rotating at room temperature, then washed for 5, 10, and 15 min at room temperature in phosphate buffer supplemented with 0.1% (w/v) Triton X-100 (PBT). After removing the wash buffer, 100 μl phosphate buffer was added and ovaries separated into ovarioles by repetitive pipetting using a P200 pipette with a tip cut to enlarge the orifice. Disrupted ovarioles were transferred to 0.2 mL tubes and incubated rotating at 4°C overnight with primary antibodies in PBT supplemented with 5% (v/v) normal donkey serum (Sigma #D9663). Primary antibodies were diluted 1:1000 for rabbit polyclonal anti-Armi C-terminal peptide (gift of William Theurkauf; Cook et al., 2004), 1:200 for mouse clone 2F8A9 monoclonal anti-Armi N-terminal peptide (1.3 mg/ml; gift of Mikiko Siomi; Saito et al., 2010, #184;), 1:500 for mouse monoclonal clone M2 anti-FLAG (Sigma), 1:200 for mouse monoclonal anti-ATP5A (Abcam 15H4C4), 1:2000 for rabbit polyclonal anti-Aub (MA514, 2.4 mg/mL, (Li et al., 2009a), 1:2000 for mouse monoclonal anti-Ago3 (gift from Julius Brennecke; Senti et al., 2015), 1:50 rat monoclonal anti-Vasa (Developmental Studies Hybridoma Bank).

The next day, ovarioles were washed three times with PBT, for 5, 10, and 15 min at room temperature. Secondary antibodies diluted in PBT were added and the ovarioles incubated in the dark, rotating at room temperature for 2 h. All secondary antibodies are from Thermo Fisher, produced in donkey or goat, against mouse, rabbit or rat IgG (H+L), highly cross-adsorbed against close species, and conjugated to Alexa Fluor 488 or Alexa Fluor 594 (for three color experiments), or Alexa Fluor 488, Alexa Fluor 546 or Alexa Fluor 633 (for four color experiments).

Ovarioles were next washed twice in PBT for 10 min each at room temperature, then incubated with 0.5 μg/mL 4’,6-diamidino-2-phenylindole diluted in 0.3 M NaCl, 0.03 M sodium citrate for 15 min at room temperature, and washed again with PBT for 10 min. Wash buffer was removed and a drop of VECTASHIELD Mounting Medium (Vector Laboratories #H-1000) added. After gentle pipetting to mix, the ovarioles in mounting medium were transferred to a glass slide using a P200 pipette with a cut tip and covered with a 0.13–0.17 mm thick cover slip (VWR #48393106). The cover slip was gently pressed with a Kimwipe to absorb extra mounting medium, then sealed with nail polish. Images were captured using a Leica TCS SP5 II laser scanning confocal microscope using a 63× HCX PL APO CS oil immersion objective (NA = 1.4) and 1 μM-thick z-stacks.

### Image Quantification

Confocal images were quantified using custom-built scripts in CellProfiler (Kamentsky et al., 2011). To measure the amount of Aub signal overlapping with Armi, primary Armi and Aub objects in separate fluorescent channels were identified using the adaptive Otsu thresholding method. “Threshold correction factor” and “lower bound on threshold” were empirically determined using representative test images. Once optimized, the same object identification settings were applied to all samples. Armi objects were then used as masking objects to mask Aub objects. The amount of signal in masked Aub objects was divided by the amount of signal in total Aub objects for each image. The amount of Vasa signal overlapping with ATP5A or Aub were quantified similarly.

### Transmission Electron Microscopy

Fly ovaries were quickly dissected in dissecting solution and transferred to 1.7 ml microfuge tubes on ice. Dissecting solution was removed and ovaries fixed in 2.5% glutaraldehyde (w/v) in 0.1 M sodium cacodylate buffer, pH 7.2, overnight at 4°C. Samples were processed and analyzed at the University of Massachusetts Medical School Electron Microscopy Core Facility using standard procedures. Briefly, intact ovaries were rinsed three times in the fixation buffer and post-fixed with 1% osmium tetroxide (w/v) for 1 h at room temperature. Samples were then washed three times with water for 10 min each and dehydrated using a graded series of 10%, 30%, 50%, 70%, 85%, and 95% ethanol, finishing with three changes in 100% ethanol. Samples were then infiltrated first with two changes 100% propylene oxide and then with a mixture of 50% propylene oxide/50%SPI-Pon 812 resin. The sample was incubated in seven successive changes of fresh 100% SPI-Pon 812 resin over 2 days, then polymerized at 68°C in flat molds. The samples were then reoriented for horizontal sections of the center of individual ovaries. Thin sections (∼70 nm) were placed on gold support grids, and stained with lead citrate and uranyl acetate to increase contrast. Sections were examined using a CM10 with 80 KV accelerating voltage, and images captured using a Gatan TEM CCD camera.

### Immunoprecipitation

***Zuc immunoprecipitation:*** freshly dissected *zucWT-3×FLAG* ovaries were crosslinked with DTME and kept on ice. For each mg of the ovary pellet, 4 μl ice-cold lysis buffer (50 mM Tris, 150 mM NaCl, 1 mM EDTA, 0.5% IGEPAL CA-630 (Sigma), 1% Empigen BB (w/v, Sigma), 0.1% SDS (w/v), 0.5 mM DTT) containing protease inhibitor cocktail (1 mM AEBSF (4-(2-aminoethyl)benzenesulfonyl fluoride hydrochloride [EMD Millipore #101500], 0.3 μM Aprotinin [Bio Basic Inc #AD0153], 20 μM Bestatin [Sigma Aldrich #B8385], 10 μM E-64 [(1S,2S)-2-(((S)-1-((4-Guanidinobutyl)amino)-4-methyl-1-oxopentan-2-yl)carbamoyl)cyclopropanecarboxylic acid; VWR #97063], and 10 μM Leupeptin [Fisher Scientific #108975]. Ovaries were homogenized with a motorized plastic pestle (Fisher Scientific #12141364) for 30 strokes on ice. The tube containing ovary lysate was then submerged in an ice-water bath and sonicated (Branson Digital Sonifier Model 450) at 40% amplitude for 2 min total sonication (8 cycles of 15 s sonication followed by 1 min rest). Lysate was centrifuged at 13,000 × *g* for 10 min at 4°C to remove insoluble material and the supernatant retained. Mouse monoclonal clone M2 anti-FLAG antibody (clone M2, Sigma) or mouse anti-GFP antibody (clone GF28R, Invitrogen, as negative control) was added at 6 μg antibody per 1 ml of lysate and incubated rotating 2 h at 4°C, before transferring the supernatant to a new tube containing protein G paramagnetic Dynabeads (1/10 volume of the lysate before removing the buffer) washed three times with lysis buffer. The tube was further rotated for 1 h at 4°C. Protein G beads were collected using a magnetic stand, washed at room temperature for 1 min successively with a volume equal to that of the original lysate of wash buffer (50 mM Tris, 1 mM EDTA, 0.05% v/v NP-40, and 0.1% w/v SDS) containing 150 mM, 300 mM, 500 mM, 750 mM and 150 mM NaCl. Beads were then washed with 0.05% v/v NP-40 in water. Finally, beads were eluted by incubating in elution buffer (50 mM Tris, 150 mM NaCl, 1 mM EDTA, 0.05% NP-40, 0.1% SDS, 1 mM DTT) containing 0.5 μg/μl (175 μM) 3×FLAG peptide (Sigma) for 10 min at room temperature with occasional mixing to keep beads suspended.

***Armi immunoprecipitation:*** Ovaries with germline-specific overexpression of Flag-Myc-tagged Armi were dissected and immunoprecipitated as described for Zuc, except that the cleared lysate was diluted by adding four volumes 50 mM Tris, 150 mM NaCl, 1 mM EDTA, 0.1% SDS, 0.5 mM DTT, and protease inhibitor cocktail. For each 1 ml diluted lysate, 6 μg mouse clone M2 monoclonal anti-FLAG antibody (Sigma) was added. An amount of Protein G Dynabead suspension equal to 1/10 volume of the diluted lysate was used.

### Western Blotting

For each mg tissue, ovaries were homogenized with a motorized plastic pestle (Thermo Fisher #12141364) in 4 μl ice-cold lysis buffer (100 mM potassium acetate, 30 mM HEPES-KOH [pH 7.4], 2 mM magnesium acetate, 1 mM DTT) containing protease inhibitor cocktail. Lysate was centrifuged at 13,000 × *g* for 10 min at 4°C and an equal volume of 2 × denaturing loading buffer (100 mM Tris-HCl [pH 6.8], 4% (w/v) SDS, 0.2% (w/v) bromophenol blue, 20% (v/v) glycerol, 200 mM DTT) was added to the supernatant and heated to 95°C for 5 min.

The lysate was resolved by electrophoresis through a 4–20% gradient polyacrylamide gel (Bio-Rad Laboratories #5671085). Proteins were transferred to a 0.45 μm pore nitrocellulose membrane (Amersham #GE10600002) in a Mini Trans-blot tank at 15 V overnight. The membrane was blocked in Blocking Buffer (Rockland Immunochemicals #MB-070) at 4°C for 5 h or overnight, then incubated overnight at 4°C in with primary antibodies diluted in Blocking Buffer. Primary antibodies were diluted 1:2000 for mouse clone 2F819 monoclonal anti-Armi N-terminal peptide (1.3 mg/ml; gift from Mikiko Siomi; Saito et al., 2010, #184;), 1:500 for goat anti-Armi C-terminal peptide (Santa Cruz dD-17), 1:10,000 for mouse clone M2 monoclonal antiFLAG (Sigma), and 1:3000 for rabbit anti-Dicer-2 (Abcam ab4732).

The membrane was washed three times, 5 min each, with TBST (50 mM Tris-HCl, pH 7.5, 150 mM NaCl, 0.1% v/v Tween 20) at room temperature, incubated for 1 h at room temperature with secondary antibodies conjugated to IRDye 680RD or 800CW (LICOR Biosciences) diluted 1:20,000 in TBST, and then washed five times, 5 min each in TBST at room temperature. All secondary antibody steps were conducted in the dark. Signal was detected using the Odyssey Infrared Imaging System.

### Mass spectrometry

FLAG immunoprecipitation was eluted as described above, and DTME was reverse-crosslinked. LC/MS/MS digestion and analysis were carried out by the University of Massachusetts Proteomics Core: the eluted immunoprecipitation reaction (for Zuc immunoprecipitation, eluate from ∼350 mg of ovary tissue; for Armi immunoprecipitation, from ∼40 mg ovary tissue) was denatured in 2× denaturing loading buffer and electrophoresed for 20 min at 100 V through an SDS-polyacrylamide gel to separate proteins from lower molecular weight contaminants. The entire protein region of the gel excised as a single band and subjected to in-gel trypsin digestion after reduction with DTT and alkylation with iodoacetamide. Peptides eluted from the gel were lyophilized and re-suspended in 25 μl 5% acetonitrile, 0.1% TFA. A 3 μl injection was loaded at 4.0 μl/min for 4 min onto a 100 μM I.D. fused-silica pre-column packed with 2 cm of 5 μM (200Å) Magic C18AQ (Bruker-Michrom) run in 5% acetonitrile, 0.1% formic acid using a NanoAcquity UPLC (Waters). Peptides were loaded onto a 75 μm I.D. gravity-pulled analytical column packed with 25 cm of 3 μm Magic C18AQ particles (100 Å), and eluted at 300 nl/min over 60 min using a 5–5% linear gradient of mobile phase B (acetonitrile + 0.1% formic acid) in mobile phase A (water + 0.1% formic acid). Ions were introduced by positive electrospray ionization via liquid junction into a Q Exactive hybrid mass spectrometer (Thermo Fisher). Mass spectra were acquired over *m/z* 300-1750 at 70,000 resolution (*m/z* 200) and data-dependent acquisition selected the top 10 most abundant precursor ions for tandem mass spectrometry by HCD fragmentation using an isolation width of 1.6 Da, maximum fill time of 110 ms, AGC target of 10^6^, collision energy of 27, and a resolution of 17,500 (*m/z* 200). For data analysis, raw data files were peak processed with Proteome Discoverer 2.1 (Thermo Fisher) followed by identification using Mascot Server 2.5 against the *Drosophila melanogaster* Uniprot FASTA file downloaded 5/2016. Search parameters included Trypsin/P specificity, up to two missed cleavages, fixed carbamidomethyl on cysteine, and the variable modifications of oxidized methionine, pyroglutamic acid for N-terminal glutamine peptides, and N-terminal acetylation of the protein. Assignments were made using a 10 ppm mass tolerance for the precursor and 0.05 Da mass tolerance for the fragments. All non-filtered search results were then loaded into Scaffold Viewer (Proteome Software, Inc.) with thresholding to a peptide FDR of 1%, for subsequent peptide/protein validation and label free quantitation.

Quantification was performed using Scaffold Viewer, considering only proteins that passed the Fisher’s exact test using weighted spectra and a threshold of Benjamini-Hochberg multiple test corrected *p* < 0.05 from three biological replicates. iBAQ enrichment score for each protein was calculated by dividing normalized iBAQ quantification value from the experimental immunoprecipitation by that from the control immunoprecipitation. A pseudo-count equal to the average of the lowest 10 iBAQ values in each sample was added to all proteins in that sample.

### Armi Structure Modeling

The predicted Armi helicase core, amino acids 692 to 1160, was submitted to I-TASSER (Roy et al., 2010). The modeled structure was superimposed on human Upf1 helicase core structure (PDB ID 2GJK) using PyMol v1.3 (Schrodinger, LLC).

### Small RNA-seq Libraries

Small RNA libraries were constructed as described (Han et al., 2015a). Briefly, total RNA (50 μg) was purified by 15% urea polyacrylamide gel electrophoresis, selecting for 18–30 nt small RNAs using 18 nt and 30 nt size markers. Half of the purified sRNAs were oxidized with 25 mM NaIO4 to deplete miRNAs and enrich for siRNAs and piRNAs (Li et al., 2009a). To reduce ligation bias, a 3’ adaptor with three random nucleotides at its 5’ end was used (5’-rApp NNN TGG AAT TCT CGG GTG CCA AGG /ddC/-3’). The 3’ adaptor was ligated to RNAs using purified, recombinant truncated, K227Q mutant T4 RNA ligase 2 at 16°C overnight, small RNA precipitated and size selected by comparison to 18 nt and 30 nt size markers ligated to the same 3’ adaptor. To exclude 2S rRNA from sequencing libraries, 10 pmol 2S blocker oligo was added before 5’ adaptor ligation (Wickersheim and Blumenstiel, 2013). The 5’ adaptor was ligated using T4 RNA ligase (Life Technologies #AM2141) at 25°C for 2 h, followed by reverse-transcription using AMV reverse transcriptase (New England Biolabs #M0277L) and PCR using AccuPrime *Pfx* DNA polymerase (Invitrogen #12344-024). PCR products were purified rom a 2% agarose gel, and the DNA recovered from the gel with QIAquick gel extraction kit (Qiagen). Length distribution and quality of the libraries were analyzed using a Bioanalyzer 2100 (Agilent). Libraries were quantified using KAPA library quantification kit (Kapa Biosystems), before being sequenced on a NextSeq500 (Illumina) to obtain 75 nt single-end reads.

The sequence of the 3’ adapter, including its 5’ three random nucleotides, was removed from raw reads. Small RNA-seq analysis was performed with piPipes 1.4 (Han et al., 2015b). Briefly, reads were aligned to rRNA and miRNA hairpin sequences using Bowtie 1.0.0 (Langmead et al., 2009). Unaligned reads were mapped to fly genome release dm3 using Bowtie and 23–29 nt piRNAs were retained for analysis. The abundance of piRNAs overlapping genes and transposons were apportioned by the number of times each piRNA aligned in the genome. Division of transposon families into germline, somatic and intermediate groups was according to Wang et al. (2015).

Ping-pong *Z_10_* (5’-to-5’ distance on opposite genomic strands = 10 nt; positions 0–9 and 11–20 as background) and phasing *Zo* (3’-to-5’ distance on the same genomic strand = 0 nt; positions 1–50 as background) scores were calculated as described (Zhang et al., 2011; Han et al., 2015a).

### RNA-seq Libraries

RNA-seq libraries were constructed as previously described (Zhang et al., 2012b) with modifications, including the use of unique molecular identifiers to eliminate PCR duplicates (Fu et al., 2018) To deplete ribosomal RNA, RNA was hybridized in 10 μl with a pool of 186 rRNA antisense oligos (0.05 μM/each) in 10 mM Tris-HCl [pH 7.4], 20 mM NaCl, heated to 95°C, cooled at –0.1°C/sec to 22°C, and then incubated at 22°C for 5 min. Thermostable RNase H (10 U, Lucigen #H39500) were added and incubated at 45°C for 30 min in 20 μl containing 50 mM Tris-HCl [pH 7.4], 100 mM NaCl, and 20 mM mgCl2. RNA was treated with 4 U Turbo DNase (Thermo Fisher #AM2238) in 50 μl at 37°C for 20 min, then purified using RNA Clean & Concentrator-5 (Zymo Research #R1016), to enrich for RNA >150 nt. RNA-seq libraries were sequenced using a NextSeq500 (Illumina) to obtain 75 + 75 nt, paired-end reads.

RNA-seq analysis was performed with piPipes. Briefly, RNAs were first aligned to rRNA sequences using Bowtie2 (v2.2.0; (Langmead and Salzberg, 2012) and unaligned reads mapped to fly genome release dm3 using STAR (v2.3.1; (Dobin et al., 2013). The numbers of reads overlapping genes and transposons were apportioned by the number of times each read aligned in the genome. Germline, somatic and intermediate transposon family classification was according to Wang et al., 2015.

### 5’ monophosphorylated long RNA Sequencing

5’ monophosphorylated long RNA-seq libraries were constructed as described (Han et al., 2015a) with modifications, including the use of unique molecular identifiers to eliminate PCR duplicates (Fu et al., 2018). PFA-crosslinked Armi immunoprecipitation eluate was reversed by adding an equal volume of 200 mM Tris-Cl, pH 7.5, 25 mM EDTA, pH 8.0, 300 mM NaCl, 2% w/v SDS containing 0.4 mg/ml proteinase K and incubating at 50°C for 1 h then 65°C for 2 h, before extracting with an equal volume of acid 5:1 phenol:chloroform pH 4.5 (AMRESCO LLC, Solon, OH, USA), and centrifuged at 20,800 × g for 5 min at room temperature. One-tenth volume 3 M sodium acetate and three volumes 100% ethanol were added to the aqueous phase and the RNA precipitated by on ice for 1 h. The precipitate was recovered by centrifugation at 20,800 × g for 15 min at 4°, washed with 70% (v/v) ethanol, air dried, and dissolved in water. rRNA depletion and RNA Clean & Concentrator-5 purification were as described for RNA-seq. A 5’-adapter with Unique Molecular Identifier (UMI) (an equimolar mix of 5’-GUU CAG AGU UCU ACA GUC CGA CGA UC (N3) CGA (N3) UAC (N3)-3’ and 5’-GUU CAG AGU UCU ACA GUC CGA CGA UC (N3) AUC (N3) AGU (N3)-3’; Fu et al., 2018) was ligated to 5’-monophosphorylated RNAs using T4 RNA ligase (Ambion) at 25°C for 2 h. The ligation reaction was purified using 1.5× volume of Ampure XP beads suspension. Reverse transcription using superScript III (Life Technologies) used a primer containing degenerate sequences at its 3’ end (5’-GCA CCC GAG AAT TCC ANN NNN NNN-3’). The reverse transcription reaction was digested with 1 μl RNase H (Ambion) at 37°C for 20 min, and purified using 1.5× volume of Ampure XP beads. Purified cDNA was amplified by PCR #1, using a pair of primers that anneal to the 5’-(5’-CTA CAC GTT CAG AGT TCT ACA GTC CGA-3’) or to 3’-(5’-GCC TTG GCA CCC GAG AAT TCC A-3’) adapter. The PCR reaction was mixed with 0.7× volume of Ampure beads, and the supernatant was transferred to a new tube containing 0.5× volume of Ampure beads (total 1.2× volume) to purify 200–400 nt PCR products. subsequently, PCR #2 was carried out using the purified products of PCR #1 and the same barcoded primer set as the small RNA library cloning protocol. PCR #2 was purified with 1.1× volume of Ampure beads. All PCR reactions used Phusion polymerase (NEB). 5’ monophosphorylated long RNA-seq libraries were sequenced using a NextSeq500 (Illumina) to obtain 75 + 75 nt, paired-end reads.

Sequences of unique molecular identifiers (UMIs; NNNCCCNNNCCCNNN) were extracted from the 5’ section of read 1. PCR duplicates were removed with umi_tools 0.5.3 using the “directional” mode (Fu et al., 2018), and reads were then analyzed using piPipes. Briefly, sequences were first aligned to rRNA using Bowtie2. Unaligned reads were then mapped using STAR to fly genome release dm3, and alignments with soft clipping of ends were removed with SAMtools 1.0.0 (Li et al., 2009b). In addition to sequencing depth, libraries were normalized using reads uniquely mapping to *flamenco*. The abundance of reads overlapping genes and transposons was apportioned by the number of times each read aligned to the genome. Division of transposon families into germline, somatic and intermediate groups was according to Wang et al. (2015).

Distance probability analyses (Han et al., 2015a) were done without considering abundance, and species uniquely mapping to *flamenco* were used to normalize libraries. To determine the significance of piRNA reads sharing 5’ ends with 5’ monophosphorylated long RNAs, the *Z_0_* score for the peak at 0 was calculated using positions 1–50 as background.

